# No benefit of subsampling in ensemble coding

**DOI:** 10.64898/2026.09.11.750959

**Authors:** Long Ni, Michael S. Landy

**Affiliations:** Department of Psychology and Center for Neural Science, New York University

**Keywords:** bounded rationality, summary representation, information integration, Bayesian inference

## Abstract

Our visual system compensates for its limited information processing capacity by creating summary representations of stimulus ensembles. A longstanding debate concerns whether summary statistics are computed by integrating information across the entire ensemble or by sampling only a subset of its items. Subsampling is intuitively appealing because 1) integrating over a small subset presumably reduces processing resource expenditure and 2) observer models employing this strategy can effectively reproduce ensemble coding performance. Here, we systematically evaluate both aspects. We first show that the apparent success of subsampling models depends on a problematic assumption: that the sensory encoding noise of each item remains constant as set size increases. This assumption entails that total coding resources, quantified as Fisher information, grow unconstrained with set size. Under the realistic assumption that total coding resources are limited, such that the encoding precision for each item decreases with increasing set size, we demonstrate that an ideal observer integrating the entire ensemble achieves the same accuracy with a strictly smaller total resource budget for every set size. We further show that this resource-constrained full-integration model provides a more parsimonious, yet equally good, account of ensemble coding behavior across multiple existing datasets as well as newly collected data. Our findings call into question the necessity of the subsampling hypothesis, as restricting integration to a subset of ensemble items confers neither a theoretical nor an explanatory benefit over resource-rational full integration.

## Introduction

Ensemble coding refers to a perceptual integration process through which the visual system extracts summary statistics, such as the mean and variance of features of interest, from an array of stimuli (see Bauer, 2015; Whitney & Yamanashi Leib, 2018, for reviews). This rapid and often automatic integration process helps alleviate the information-processing bottleneck of the human visual system in the face of a continuous influx of sensory input (Allik, Toom, Raidvee, Averin, & Kreegipuu, 2014; Alvarez, 2011; Ariely, 2001; Chong & Treisman, 2003; Dubé & Sekuler, 2015).

Extensive empirical work has demonstrated the remarkable precision of ensemble coding across a wide range of lowand high-level visual feature domains (Ariely, 2001; Chong, Joo, Emmmanouil, & Treisman, 2008; Chong & Treisman, 2003, 2005a, 2005b; Dakin, 2001; Florey, Clifford, Dakin, & Mareschal, 2016; Florey, Dakin, & Mareschal, 2017; Hubert-Wallander & Boynton, 2015; Khayat & Hochstein, 2018). Observers can rapidly and seemingly effortlessly extract summary statistics from stimulus ensembles of varying set sizes, even when they are not explicitly instructed to do so (Brady & Alvarez, 2011; Hansmann-Roth, Kristjánsson, Whitney, & Chetverikov, 2021). Despite a rich characterization of ensemble coding behaviors, considerably less is known about the underlying computational mechanisms. A prominent unresolved question concerns how summary statistics are actually computed. For example, when estimating the ensemble mean, does the visual system integrate information from all items in the ensemble, a full-integration strategy, or does it rely on only a sampled subset of items, a subsampling strategy?

The subsampling hypothesis is intuitively appealing because averaging only a subset of ensemble items presumably requires fewer processing resources than integrating the entire stimulus array. This proposal has been motivated partly by evidence that focused attention and visual working memory can select or maintain only a small number of individuated objects at a time (Alvarez & Cavanagh, 2005; Cowan, 2001; Luck & Vogel, 1997; Pylyshyn & Storm, 1988). On this account, items that fall outside these capacity-limited representations would contribute little or nothing to the computed ensemble statistic (Allik, Toom, Raidvee, Averin, & Kreegipuu, 2013; Myczek & Simons, 2008; Zepp & Dubé, 2026). What makes the subsampling hypothesis perhaps even more attractive is the finding that integrating only a small subset of ensemble items appears sufficient to account for human ensemble-coding performance across a wide range of stimulus features (Allik et al., 2013; Dakin, 2001; Florey et al., 2016; Haberman & Whitney, 2010; Im & Halberda, 2013; Marchant, Simons, & de Fockert, 2013; Maule & Franklin, 2016; Myczek & Simons, 2008; Solomon, May, & Tyler, 2016; Sweeny, Wurnitsch, Gopnik, & Whitney, 2015; Tokita, Ueda, & Ishiguchi, 2016; Virtanen, Olkkonen, & Saarela, 2020; Zepp & Dubé, 2026). In these subsampling models, the effective sample size, the number of ensemble items required to be integrated to match observed performance, is typically much smaller than the actual ensemble size, *N*. Although early claims that ensemble coding requires the integration of only one to three items have largely been rejected, a commonly adopted account assumes that the effective sample size increases approximately in proportion to 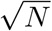 (Whitney & Yamanashi Leib, 2018).

There are, however, several problems with the subsampling hypothesis. First and foremost, the apparent success of subsampling models in accounting for human performance is contingent on a problematic assumption: that the sensory encoding noise associated with each individual item remains constant as the effective sample size increases. Previous studies testing subsampling models have either omitted sensory encoding noise altogether (Marchant et al., 2013; Myczek & Simons, 2008) or assumed that it remains constant across effective sample sizes (Allik et al., 2013; Florey et al., 2016; Haberman & Whitney, 2010; Im & Halberda, 2013; Maule & Franklin, 2016; Sweeny et al., 2015; Tokita et al., 2016; Virtanen et al., 2020; Zepp & Dubé, 2026). This assumption implies that the total processing resources, quantified by the summed encoding precision across items, is unconstrained and increases linearly with the effective sample size. Extensive evidence instead indicates that the encoding precision of individual stimuli decreases monotonically as a limited pool of processing resources is distributed across an increasing number of items (Alvarez & Franconeri, 2007; Bays, Catalao, & Husain, 2009; Keshvari, Van den Berg, & Ma, 2013; Ma & Huang, 2009; Palmer, 1990, 1994; Sims, Jacobs, & Knill, 2012; Tokita et al., 2016; Van den Berg, Shin, Chou, George, & Ma, 2012). This decline in item-level encoding precision is observed whether the number of items falls within or exceeds the capacity of focused attention.

A second problem with the subsampling account is its excessive flexibility (Ariely, 2008). There is no principled rule specifying how many items should be sampled, and the inferred effective sample size appears to depend on multiple factors, including the physical set size and the variance of the ensemble feature values (Dakin, 2001; Dakin, Mareschal, & Bex, 2005; Im & Halberda, 2013; Marchant et al., 2013; Maule & Franklin, 2016). Consider the effect of set size. Although the overall relationship between effective sample size and physical set size approximately follows a power law (Dakin, 2001; Whitney & Yamanashi Leib, 2018), in some studies the fitted effective sample size remains largely unchanged as the physical set size increases (Allik et al., 2013). In general, subsampling models are often overparameterized (i.e., a separate parameter indicating the effective sample size for each set size). As a result, the inferred effective sample sizes can become an ad hoc set of model parameters that are difficult, if not impossible, to verify experimentally.

A third major challenge for the subsampling account concerns its neural implementation. In most ensemble-averaging tasks, the display consists of spatially separated objects that generate distinct retinal signals and are therefore likely to be represented, at least initially, by the early visual system. It is therefore unclear at what stage of the visual-processing hierarchy subsampling is assumed to occur. One possibility is early selection, whereby only a subset of items is encoded with sufficient fidelity to contribute to the ensemble estimate. Another is late selection, whereby all items are initially encoded but only a subset of their sensory representations is granted access to downstream decoding. These alternatives require distinct neural mechanisms: the former must explain how encoding is selectively prevented or strongly attenuated for some items, whereas the latter must explain how otherwise available sensory representations are excluded from the readout. We are not aware of any subsampling accounts that specify these mechanisms, nor is there direct empirical evidence that ensemble processing involves either selective encoding failure or discrete gating of a small subset at the decoding stage. By contrast, full integration has a straight-forward neural implementation in population-response models, in which neural activity elicited by all items in the array is pooled to derive an ensemble estimate (Robinson & Brady, 2023; Utochkin, Choi, & Chong, 2023).

Here, we advocate an optimal full-integration model operating under resource constraints. It is well established that sensory systems must process information with limited computational resources. A growing body of work has therefore sought to characterize perceptual and cognitive behavior as optimal adaptation to such constraints, an approach broadly categorized as resource-rational analysis (Bhui, Lai, & Gershman, 2021; Lieder & Griffiths, 2020; Van den Berg & Ma, 2018). In the context of ensemble coding, the availability of limited processing resources that are distributed over the entire stimulus array naturally implies that sensory encoding noise increases with set size. Within this resource-rational framework, we evaluate two key aspects of the subsampling account. Specifically, we ask 1) whether a subsampling strategy, often assumed to operate within focused attention, is more resource efficient compared to the resource-constrained full-integration model, and 2) whether subsampling provides a sufficient and superior account of ensemble-coding behavior?

Our systematic analyses reject both claims. We first show, perhaps counterintuitively, that a resource-constrained optimal full-integration model can achieve the same level of ensemble-coding accuracy with a strictly lower coding-resource budget—quantified as Fisher information—than a subsampling model for any set size (*N >* 1). Equivalently, for a fixed resource budget, distributing processing resources across the entire stimulus array always yields greater ensemble averaging accuracy than concentrating resources on a subset of items. This result provides a normative basis for empirical findings that distributed attention improves ensemble-coding performance relative to focused attention (Baijal, Nakatani, van Leeuwen, & Srinivasan, 2013; Chong et al., 2008; Chong & Treisman, 2003, 2005a, 2005b). We then evaluate the two models against multiple existing and newly collected datasets and show that the full-integration model accounts at least as well for ensemble-averaging behavior across a range of stimulus features, while providing a consistently more parsimonious explanation of the data.

Our primary goal here is not to categorically rule out subsampling in ensemble coding, as the flexibility and lack of principled constraints in many subsampling models make them difficult to falsify. Rather, we aim to demonstrate that invoking subsampling is neither necessary nor advantageous: it offers no identifiable benefit over a more parsimonious full-integration account. Across our theoretical analyses and model comparisons, subsampling confers no advantage in explaining ensemble-coding behavior. It uses processing resources less efficiently and does not provide a superior account of the empirical data, despite requiring greater model flexibility. These findings are consistent with recent work showing that the visual system integrates information across ensemble stimuli in a manner that maximizes decision accuracy under resource constraints (Ni & Stocker, 2023, 2024, 2026). We therefore argue that resource-constrained full integration should serve as the default framework for modeling and interpreting ensemble-coding behavior.

This paper is structured as follows. We first formulate the resource-constrained full-integration model and analytically compare its resource efficiency with that of the subsampling model. We then evaluate the empirical efficacy of both models by fitting them to multiple datasets. Finally, we discuss the results and their implications for our understanding of ensemble coding.

## Model formulation

### Full-integration model

In this section, we formulate a resource-constrained optimal full-integration model for an ensemble-averaging task. In a typical ensemble-averaging task, the observer is presented with a stimulus ensemble Θ, with a generative mean *µ* and a standard deviation *σ*_stim_. The stimulus ensemble Θ = (*θ*_1_, *θ*_2_, …, *θ*_*N*_) consists of *N* individual items. However, the observer only has access to the noisy measurement of each individual item *x*_*i*_, *i* ∈ [1, 2, ‥, *N*] and therefore needs to infer the generative mean of the stimulus ensemble based on the measurement set *X* = (*x*_1_, *x*_2_, …, *x*_*N*_). Below, we first specify the resource-constrained encoding part of the model, followed by the optimal decoding.

### Resource-constrained encoding

The full-integration model assumes homogeneous sensory encoding, such that each item (*θ*_*i*_) is encoded with the same sensory noise level. Specifically, the noisy measurement (*x*_*i*_) is assumed to follow a Gaussian distribution centered on the true stimulus value (*θ*_*i*_) with standard deviation *σ*_sens_. Sensory uncertainty for circular features, such as orientation and hue, is often modeled using a von Mises distribution. In the high-concentration regime, however, the von Mises distribution is locally well approximated by a Gaussian. Because the circular feature values in all datasets analyzed here span only a restricted range away from the wrap-around boundary, and because both stimulus dispersion and sensory uncertainty are small relative to the relevant feature period (180 deg for orientation and 360 deg for direction or hue), we model sensory uncertainty as additive Gaussian noise for simplicity:

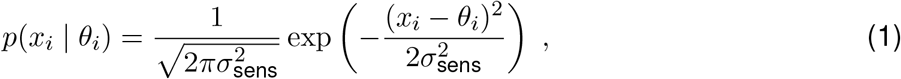

where *x*_*i*_ is the noisy measurement of each stimulus *θ*_*i*_, *i* ∈ [1, 2, …, *N*].

The model further assumes that the total coding resource is limited and distributed across all individual items in the ensemble. Consequently, the sensory encoding noise per item, *σ*_sens_, increases with set size *N*, with the relation specified as:

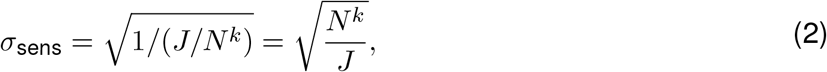

in which *J*, the Fisher information, indicates the baseline available coding resource (i.e., when *N* = 1), and *k* is the parameter of the power law. We introduce the power-law scaling to accommodate evidence that the total amount of coding resources allocated across all items does not necessarily remain fixed but may decrease or increase monotonically with set size across various tasks (see Bays et al., 2009; Bays & Husain, 2008; Elmore et al., 2011; Keshvari et al., 2013; Van den Berg et al., 2012). Van den Berg and Ma (2018) have recently shown that a normative tradeoff between behavioral performance and neural cost may underlie such power-law scaling (see Discussion).

The total Fisher information (or processing resources) *J*_t_ for a set size *N* is then expressed as:

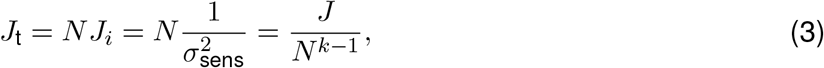

in which *J*_*i*_ denotes the Fisher information for each individual item (Fig.1a). Note that *J*_t_ equals *J* and remains constant as a function of set size when *k* = 1 (Fig.1b, black line). *J*_t_ monotonically increases and decreases with set size when *k <* 1 and *k >* 1, respectively. The resulting sensory encoding noise, *σ*_sens_, increases with set size for any *k >* 0 (Fig.1c).

**Figure 1.**
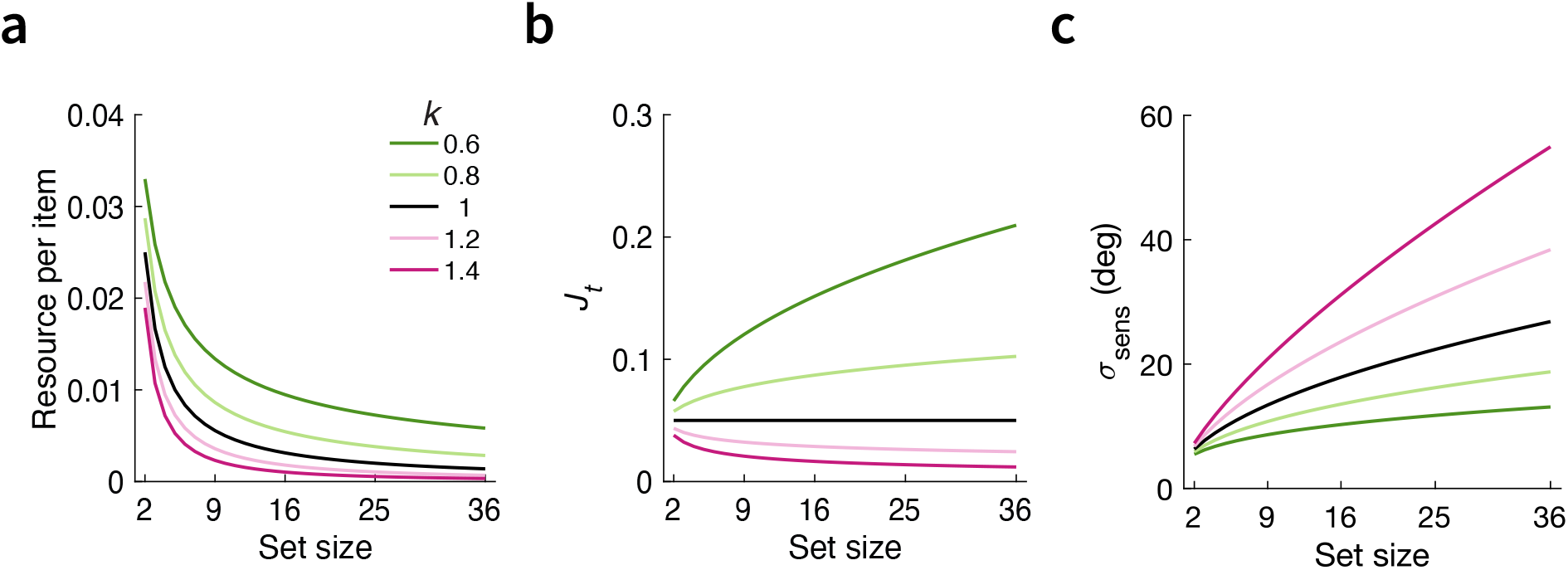
(a) Processing resource, quantified as Fisher information, per item, (b) total Fisher information, and (c) sensory encoding noise as a function of set size for different values of the power-law scaling factor *k*. All values are computed under a fixed baseline resource *J* = 0.05.

### Optimal decoding

The optimal decision stage involves inference under the hierarchical generative model, which reflects the statistical structure of an ensemble averaging task. We assume that on every trial, the ensemble stimuli are sampled from a Gaussian distribution with a generative mean *µ* and a standard deviation *σ*_stim_. Given the measurement set *X* of the stimulus ensemble Θ, the observer has to infer *µ* by computing the posterior distribution (see Appendix A for derivation):

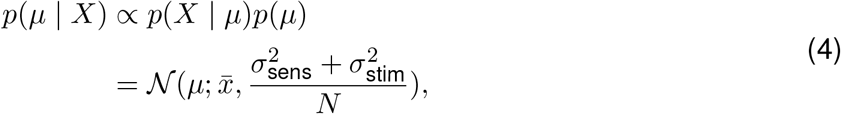

in which 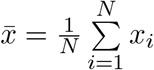 and the prior over *µ, p*(*µ*), is assumed to be uniform.

The distribution of the estimated ensemble mean 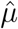 given one particular stimulus set Θ is then computed by marginalizing over the measurement of each stimulus (see Appendix B for derivation):

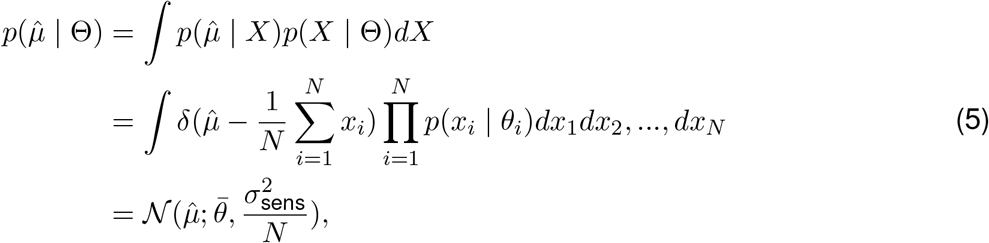

in which 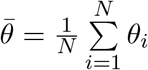.

Finally, we can compute the distribution of 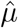 conditional on the true generative mean (*µ*), rather than on a particular stimulus set (Θ) presented on a given trial.

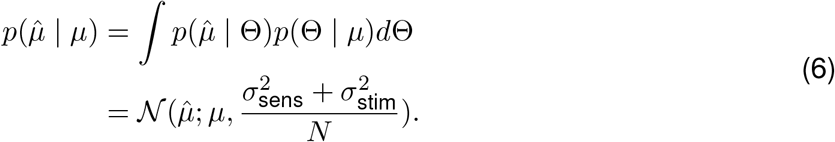

Notably, the variance of the estimated ensemble means 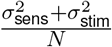 in Eq. 6 takes a similar form to the equivalent-noise equation used to estimate internal noise and “averaging” (or “sampling”) efficiency with *N* being treated as a free parameter (i.e., the number of samples; Dakin, 2001; Dakin et al., 2005; Solomon, 2010). Here we derived Eq. 6 under the assumption of Gaussian stimulus variability and additive Gaussian sensory noise.

So far, we have derived the optimal response for an ensemble-averaging estimation task. We now consider the corresponding behavior in an ensemble-discrimination task (e.g., deciding which of two stimulus ensembles has the larger mean). The probability of reporting the test ensemble as having the larger mean is given by

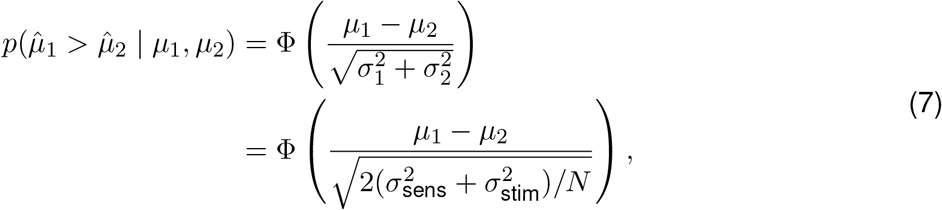

in which 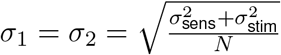, and Φ is the standard normal CDF.

### Subsampling model

The subsampling model tested here shares the same optimal decoding rule as the full-integration model but differs in its encoding process. Specifically, following typical subsampling accounts, we assume that the observer computes the summary statistics from a randomly selected subset of *M* ensemble items (*M < N*) and that the sensory encoding noise for each sampled item remains constant as *M* increases (Allik et al., 2013; Florey et al., 2016, 2017; Myczek & Simons, 2008; Simons & Myczek, 2008). Under this model, the total coding-resource expenditure is given by

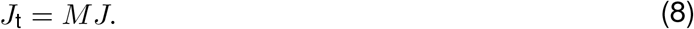

Note that our formulations of the full-integration and subsampling models include only their core computational components. This simplification allows differences in resource expenditure and behavioral predictions to be attributed directly to their distinct encoding strategies, the key computational distinction examined here. This minimal formulation, which already provides a good account of the data as shown below, makes the comparison more general and less dependent on assumptions about secondary processes. Additional components, such as decision noise or explicit computational costs, could be incorporated without altering our main conclusions, provided they are applied comparably to both models.

## Results

### Full integration is more resource-rational

With the two models formally specified, we first ask whether subsampling is more resource-efficient than full integration. To address this question, we compare the total resource expenditure required under each model to achieve the same level of ensemble-averaging accuracy. As a first step, we derive the number of sampled items, *M*, required by the subsampling model to match the performance of the full-integration model. For an ensemble-averaging estimation task, matched performance corresponds to equal variance of the resulting estimates. Therefore, we have

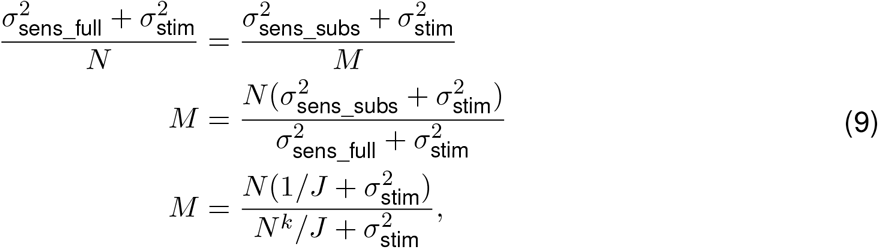

where *σ*_sens_full_ and *σ*_sens_subs_ are sensory encoding noise under the full-integration model and the sub-sampling model, respectively. As shown in Fig. 2a, to match the performance of the full-integration model, the number of stimuli required to be sampled (*M*) depends strongly on *k. M* increases with physical set size when *k* ≤ 1, but the rate of this increase is higher at smaller *k* values. When *k >* 1, *M* can even decrease with physical set size, indicating a non-monotonic relationship. The dependence of *M* on *k* and *J* specified in Eq. 9 can be more clearly characterized by examining the derivative of *M* with respect to *N*.

**Figure 2.**
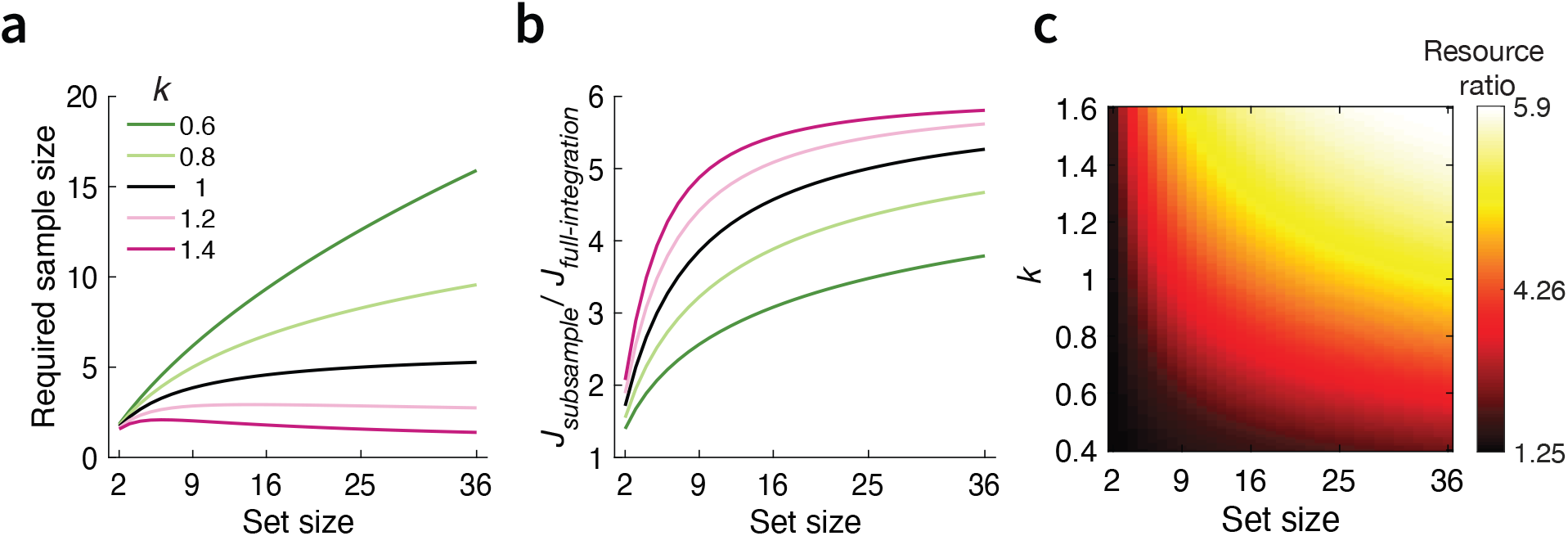
**(a)** Required sample size under the subsampling model to match the performance of the full-integration model, plotted as a function of physical set size for different values of the power-law scaling parameter *k*. **(b)** Ratio of total Fisher information under the subsampling and full-integration models, matched for performance, as a function of physical set size for different values of *k*. **(c)** Heatmap of the same Fisher-information ratio as a joint function of physical set size and *k*. All values are computed under a fixed baseline resource *J* = 0.05 and external stimulus noise *σ*_stim_ = 10.

Next, we can compute the total amount of coding resource expenditure under subsampling,

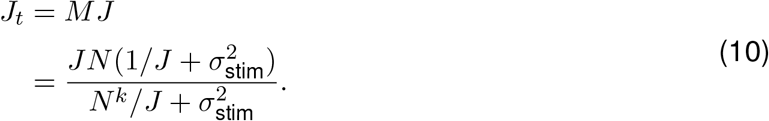

Finally, we compare the total resource expenditure under the two models when they achieve the same accuracy by computing the ratio between 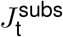 and 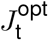.

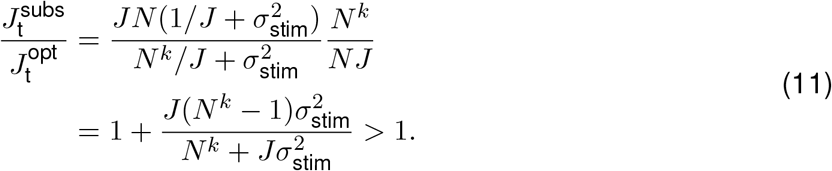

Eq. 11 indicates that subsampling always requires greater coding resources to achieve the same level of performance as the full-integration model for any set size *N >* 1 and *k >* 0. Moreover, the ratio between the required resource expenditures under the two models monotonically increases with set size *N*, the power-law scaling factor *k* (see example in Fig. 2b and c), and the baseline available coding resource *J*. These analytical results show that subsampling is far less resource-efficient compared to full integration.

So far, we have derived analytical solutions under the assumptions that ensemble items are drawn from a Gaussian distribution and that the observer estimates the generative mean. Neither assumption is necessary for our main conclusion. To achieve the same level of ensemble coding accuracy, subsampling consistently requires greater coding resources when ensemble items are drawn from a uniform distribution (Appendix C) and when the observer instead estimates the sample mean (Appendix D).

### Full integration offers a parsimonious account of set size effects

Ensemble set size has been shown to modulate ensemble coding accuracy. Many studies have reported improved performance, such as lower discrimination thresholds or reduced estimation variance, as set size increases (Baek & Chong, 2020; Chong et al., 2008; Haberman & Whitney, 2010; Lee, Dague, Sobel, Paternoster, & Puri, 2021; Robitaille & Harris, 2011; Solomon, Morgan, & Chubb, 2011; Virtanen et al., 2020), whereas others have found relatively stable performance across set sizes (Allik et al., 2013; Ariely, 2001; Chong & Treisman, 2005b; Maule & Franklin, 2015). Under subsampling accounts, such set size effects are often accommodated rather than predicted: a separate effective sample size parameter is typically estimated for each physical set size, allowing the inferred number of integrated items to vary across conditions.

In contrast, the full-integration model predicts how ensemble coding accuracy varies with set size given the scaling of total resource expenditure and the degree of stimulus-feature variability. To illustrate this, we compute the variance of the ensemble-mean estimates as a joint function of set size *N* and the power-law scaling parameter *k* (see Eq.6):

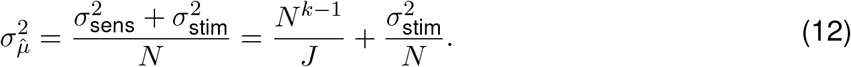

Its derivative with respect to *N* is

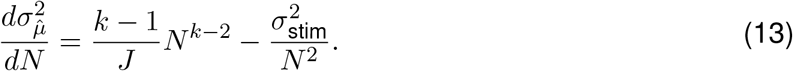

Eq. 13 indicates that changes in ensemble coding precision across set sizes depend strongly on *k*:

- *k <* 1 (Increasing resource allocation): Total resource expenditure grows with *N*, causing 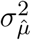 to decrease monotonically with set size 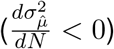 (Fig. 3, bottom panels, green lines).
- *k* = 1 (Constant resource allocation): Total resource expenditure remains fixed across set size. 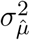 still decreases with set size 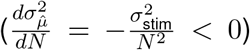, but approaches an asymptotic variance floor determined by sensory encoding noise 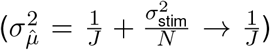 (Fig. 3, bottom panels, black line).
- *k >* 1 (Decreasing resource allocation): The relationship between 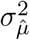 and *N* is no longer monotonic, as the two terms in Eq. 12 move in opposite directions. Sensory noise (*N* ^*k*−1^*/J*) term increases with *N*, whereas stimulus variability 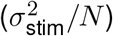 term decreases with 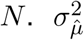 thus decreases at smaller set sizes where stimulus variability dominates, but increases at larger set sizes where sensory noise dominates (Fig. 3, bottom panel<u>s, pin</u>k lines). This U-shaped transition reaches its minimum at the critical set size 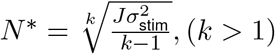.

**Figure 3.**
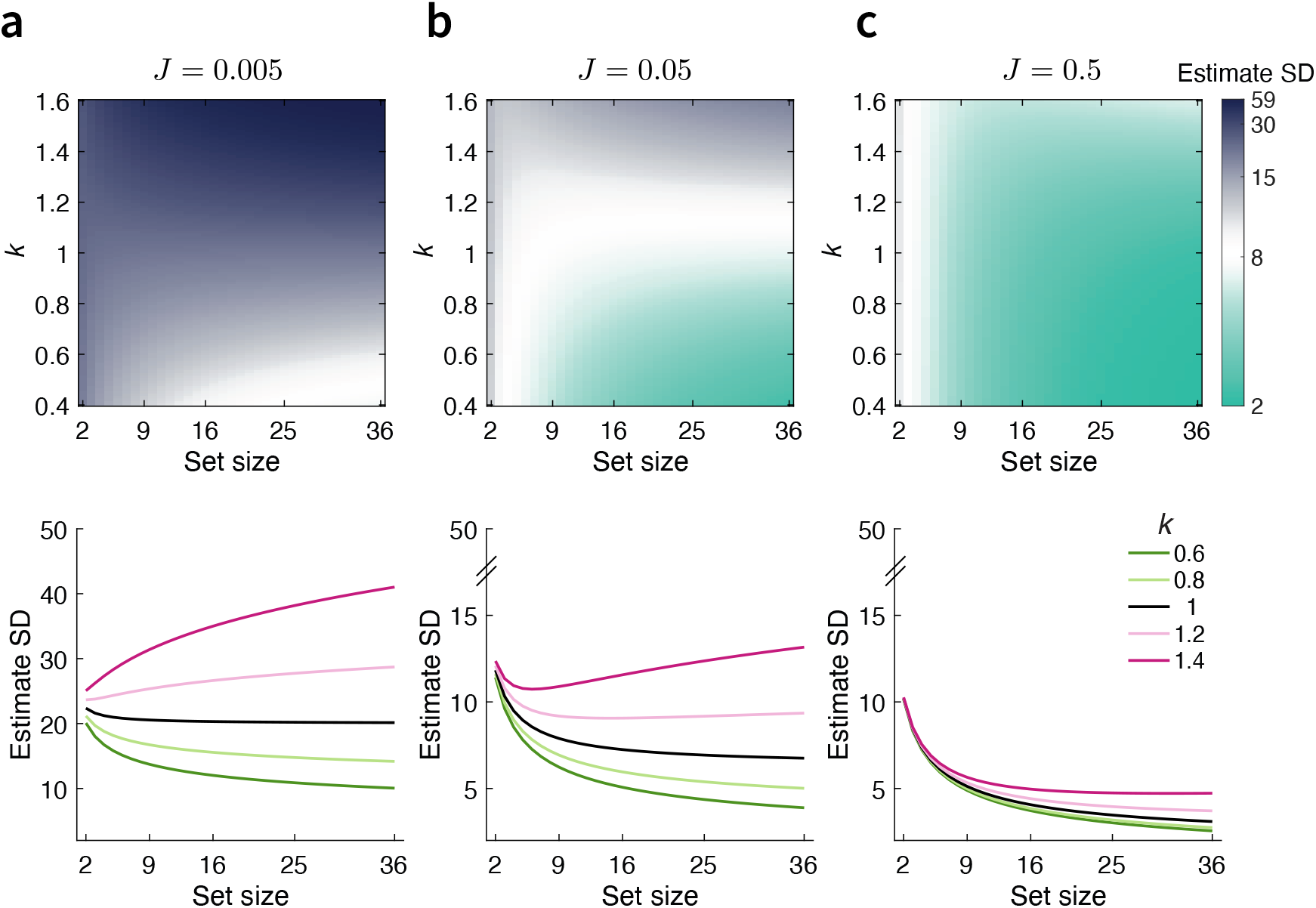
Predicted ensemble-mean estimation performance across set sizes and resource scaling parameters. **(a–c)** Standard deviation (SD) of ensemble-mean estimates shown as a joint function of physical set size *N* and power-law scaling parameter *k*, computed for baseline coding resource levels *J* = 0.005, 0.05, and 0.5, respectively. Bottom panels display 1D slices through the corresponding top heatmaps for selected values of *k*. Stimulus-feature variability is fixed at *σ*_stim_ = 10.

Under low baseline resource levels (e.g., *J* = 0.005), this U-shaped transition occurs only at a relatively small critical set size *N* ^∗^. Consequently, when *k >* 1, ensemble coding performance overall degrades (i.e., yielding a higher standard deviation of ensemble-mean estimates) as set size increases (Fig. 3a). A higher baseline coding resource (e.g., *J* = 0.05), which reduces sensory encoding noise, shifts *N* ^∗^ toward larger set sizes (Fig. 3b). If *J* increases further (e.g., *J* = 0.5) such that sensory noise becomes negligible relative to stimulus variability across the experimentally relevant range, estimation variance continues to decrease with set size (Fig. 3c).

The full-integration model therefore offers a more parsimonious account of set-size effects: changes in the allocation of processing resources determine how ensemble-averaging performance varies with set size. As the amount of sensory input increases substantially with set size, the visual system could reasonably increase its total resource allocation or maintain it at a comparable level. This provides a natural explanation for why ensemble-averaging performance often improves with increasing set size. Improved performance with increasing set size can also arise when baseline coding resources are sufficiently high, although this may represent a less typical regime.

### Empirical evaluation of the models

Our mathematical analyses have shown that, once processing-resource constraints are taken into account, the subsampling account offers no theoretical advantage over full integration. It consistently requires greater coding resources to achieve the same level of ensemble-coding precision and provides no principled account of set-size effects in ensemble-averaging behavior.

We next turn to the second aspect of subsampling and ask whether it provides greater explanatory power than full integration. Much of the appeal of subsampling models stems from their ability to reproduce observed ensemble-averaging performance. To evaluate this aspect, we compare the two models against two existing datasets and one newly collected dataset spanning multiple feature domains. Across all datasets, model comparisons show that the subsampling model does not provide a superior account of behavior relative to the full-integration model.

As noted above, here we compare the simplest formulations of the two models, each including only its core computational parameters. The full-integration model has two parameters (*J* and *k*), whereas the subsampling model has *Q* + 1 parameters, with *Q* being the number of set sizes tested in the experiment. In addition to the nonparametric subsampling model typically used in the literature, we also tested a parametric variant. This variant imposes a power-law relationship between physical set size *N* and effective sample size, as summarized by previous work (Whitney & Yamanashi Leib, 2018):

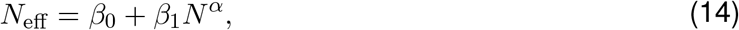

where *β*_0_, *β*_1_, and *α* are free parameters. Compared with the standard nonparametric subsampling model, this formulation constrains how effective sample size varies across set sizes and therefore reduces model flexibility and the risk of overfitting, providing a more balanced comparison with the full-integration account.

We fit each model by minimizing the negative log likelihood using Bayesian Adaptive Direct Search (Acerbi & Ma, 2017). Model performance was then compared using the Akaike Information Criterion (AIC) and Bayesian Information Criterion (BIC), both of which balance goodness of fit against model complexity.

### Ensemble averaging of hues

The first dataset we used to adjudicate the two models was from Virtanen et al. (2020), in which the authors measured performance of ensemble hue averaging across multiple set sizes. In their discrimination task, eight participants viewed two sequentially presented arrays of hue stimuli and judged whether the second array was yellower or bluer than the first (see Experiment 1 of Virtanen et al., 2020, for details)(Fig. 4a). On each trial, the hues within each ensemble were either identical (i.e., homogeneous) or randomly drawn from a von Mises distribution with low (*κ* = 40) or high dispersion (*κ* = 15). Under the Gaussian approximation, the low and high external-noise conditions correspond to standard deviations of approximately 9.12 and 14.79 degrees, respectively, defined along the hue circle in CIELAB color space. Four set sizes were tested: 1, 4, 16, and 64 (Fig. 4a). For our purposes, we excluded the homogeneous-hue conditions and the set-size-1 condition, because neither requires averaging across heterogeneous ensemble items and thus does not constitute an ensemble-averaging computation.

**Figure 4.**
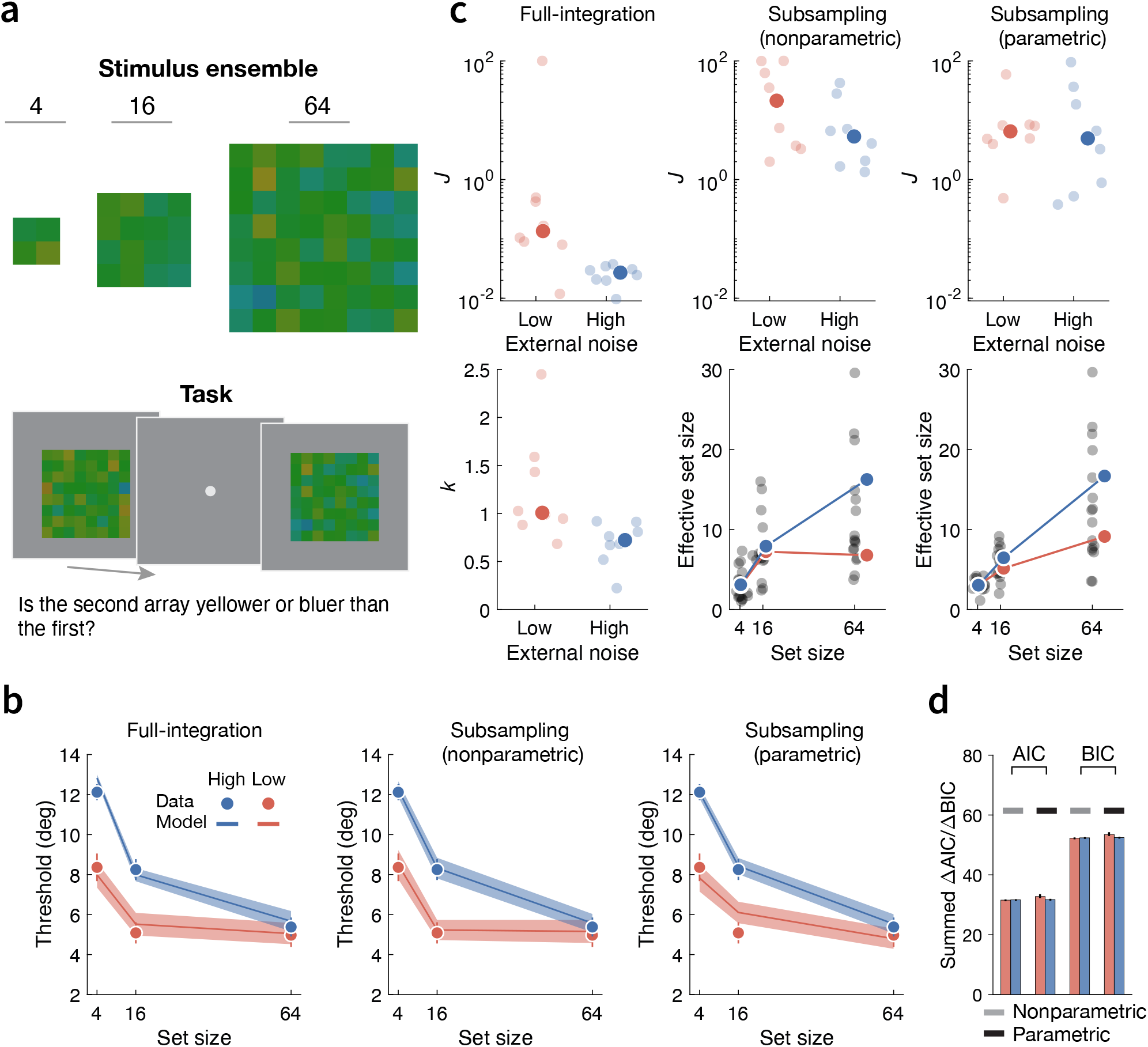
Model fits to data from Experiment 1 of Virtanen et al. (2020). (**a**) Example stimulus ensembles (top) and the ensemble mean discrimination task (bottom) used in the experiment, reproduced from Fig. 3 in Virtanen et al. (2020). (**b**) Measured and model-predicted thresholds for discriminating ensemble mean hue as a function of set size under low (red) and high (blue) external-noise conditions. Error bars and shaded regions indicate SEMs of the measured data and model predictions, respectively. (**c**) Fitted model parameters. The top row shows the fitted baseline resource parameter *J* for the full-integration (first column), nonparametric subsampling (second), and parametric subsampling (third) models. The bottom row shows the fitted scaling parameter *k* for the full-integration model (first column) and the recovered effective sample sizes for the nonparametric (second) and parametric (third) subsampling models. Dimmed dots indicate individual-participant estimates, and solid dots indicate the group median for *J* and *k* and the group mean for effective sample size. (**d**) Model comparison. Summed AIC and BIC differences between the full-integration model and each subsampling model for both external-noise conditions. Positive values favor the full-integration model. Error bars indicate 95% confidence interval over the sums obtained from 10^3^ bootstrapped samples.

We fit both the full-integration and subsampling models separately to each external-noise condition. Within each condition, the models were fit jointly to the psychometric curves across all three set sizes. Model fitting was performed for each individual participant separately. We then extracted the discrimination threshold, defined as the standard deviation of the psychometric function, from both the observed and predicted psychometric curves.

The full-integration model and the two subsampling models closely captured the measured discrimination thresholds across all set sizes in both external-noise conditions (Fig. 4). Consistent with the data, all three models predicted that the threshold for discriminating ensemble mean hue decreased with increasing set size (Fig. 4b; see Fig. A1 for model fits to data of individual participants).

The fit parameters reveal the difference in resource allocations under the two accounts. To achieve roughly the same performance, the full-integration model required a much smaller baseline coding resource (i.e., smaller *J* value) in both external-noise conditions than either subsampling model (Fig. 4c, top row). In the full-integration model, the fitted scaling parameter *k*, which determines how the total allocation of processing resources changes with set size, was close to 1 in the low-noise condition, indicating relatively stable total resource expenditure, but fell below 1 in the high-noise condition, indicating increasing total resource expenditure with set size (Fig. 4c, bottom row, leftmost column). This difference between the two external-noise conditions may reflect flexible deployment of processing resources according to task demands: a higher information processing load under the high external-noise condition may demand greater resource allocation as set size increases. Under both subsampling models, the recovered effective sample size was substantially smaller than the physical set size in both external-noise conditions, in line with previous studies. The effective sample size was also overall larger in the high external-noise condition, with the difference between the two conditions growing with set size (Fig. 4c, bottom row, middle and rightmost columns).

Importantly, although the full-integration and subsampling models provided comparably good fits to the behavioral data, the full-integration model achieved this with a lower total resource expenditure and fewer free parameters. To compare the resource-efficiency, we computed the ratio of the actual total resource expenditure under the full-integration and the non-parametric subsampling models given their corresponding fit parameters. Consistent with our mathematical analyses, the total resource expenditure required under the subsampling model is substantially greater than under the full-integration model across set sizes (Fig. 7a, left). Its advantage in parsimony was confirmed by both AIC and BIC, which favored the full-integration model over the subsampling models in both external-noise conditions (Fig. 4d).

### Ensemble averaging of size

Next, we evaluated the models against the ensemble size averaging dataset from Baek and Chong (2020). To our knowledge, this is the only previous study that directly compared a subsampling account with a heuristic full-integration account of ensemble coding. Similar to our model assumption, their full-integration model, termed the zoom-lens model, assumes that limited attentional resources are distributed across all ensemble items, causing the encoding precision of individual items to decrease as set size increases. However, the relationship between set size and item-level sensory noise is specified using a more elaborate function.^1^ Baek and Chong (2020) measured performance in a discrimination task in which four participants viewed two sequentially presented arrays of circles varying in size and judged which array had the larger average size (Fig. 5a). Six set sizes (i.e., the number of circles in each array) were tested: 1, 2, 4, 8, 16, and 32 (see Experiment 1 of Baek & Chong, 2020, for details).

**Figure 5.**
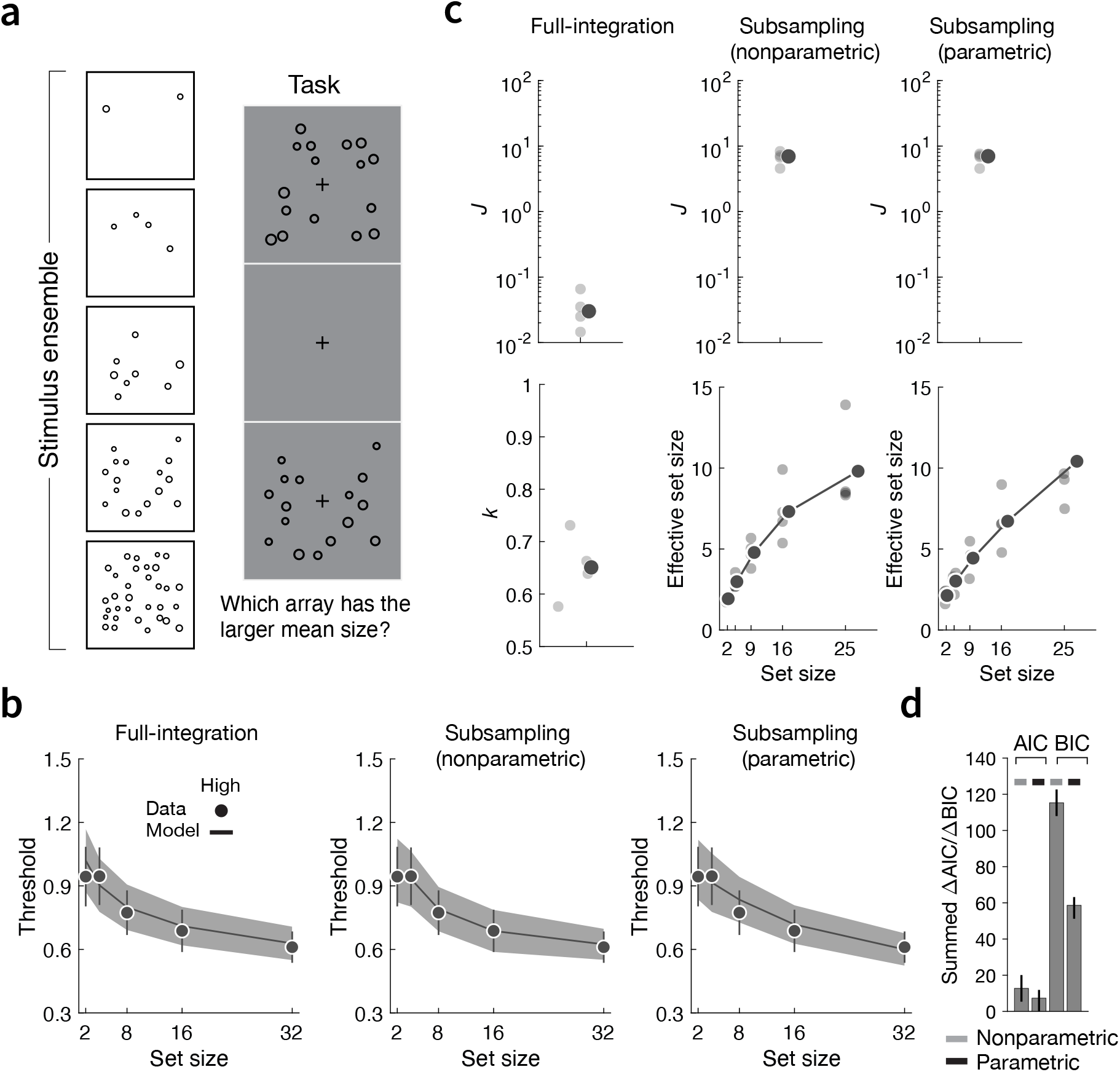
Model fits to data from Experiment 1 of Baek and Chong (2020). (**a**) Example stimulus ensembles (top) and the ensemble mean discrimination task (bottom) used in the experiment. (**b**) Measured and model-predicted thresholds for discriminating ensemble mean size as a function of set size. Error bars and shaded regions indicate SEMs of the measured data and model predictions, respectively. (**c**) Fitted model parameters. The top row shows the fitted baseline resource parameter *J* for the full-integration (first column), nonparametric subsampling (second), and parametric subsampling (third) models. The bottom row shows the fitted scaling parameter *k* for the full-integration model (first column) and the recovered effective sample sizes for the nonparametric (second) and parametric (third) subsampling models. Dimmed dots indicate individual-participant estimates, and solid dots indicate the group median for *J* and *k* and the group mean for effective sample size. (**d**) Model comparison. Summed AIC and BIC differences between the full-integration model and each subsampling model for both external-noise conditions. Positive values favor the full-integration model.

Unlike the previous dataset, for which trial-by-trial stimulus ensembles were unavailable, this dataset included the stimulus values presented on individual trials. We therefore fit the full-integration model and the two subsampling models jointly to each participant’s binary responses across all set sizes except *N* = 1, which was excluded for the same reason described above. Because the generative distribution from which individual circle sizes were sampled was not specified in the original study, we assumed a uniform distribution over stimulus sizes for all models.

As shown in Fig. 5b, the full-integration model and both subsampling models accurately reproduced the measured discrimination thresholds across all set sizes (see Fig. A2 for model fits to data of individual participants). As in the ensemble hue averaging dataset, discrimination thresholds generally decreased with increasing set size. The fitted baseline resource parameter *J* values were again substantially smaller under the full-integration model than under either subsampling model (Fig. 5c, top row). Under the full-integration model, the fitted scaling parameter *k* values were well below 1, indicating that the total allocation of coding resources increased with set size (Fig. 5c, bottom row, first column). Under both subsampling models, the recovered effective sample size increased with physical set size but remained consistently smaller than the actual set size (Fig. 5c, bottom row, second and third columns).

Given the fitted parameter values under each model, we can once again show that the subsampling models required substantially greater total coding resources than the full-integration model to achieve comparable ensemble-coding performance (Fig. 7a, middle). Both model comparison metrics further confirmed the parsimony advantage of the full-integration model (Fig. 5d). Finally, for this dataset, we also compared our optimal full-integration model with the heuristic full-integration model proposed by Baek and Chong (2020). Our model provided a consistently better account of the data across participants (Fig. A2b).

### Ensemble averaging of orientation

We have so far evaluated the models against two existing datasets that measured ensemble averaging of stimulus hue and size in discrimination tasks. In this section, we assess the full-integration and subsampling models using a new experiment in which participants estimated the average orientation of stimulus ensembles. We tested six set sizes: 2, 4, 9, 16, 25, and 36. On each trial, ensemble orientations were randomly drawn from a Gaussian distribution with a mean randomly selected between -90 and 90 deg. We tested two external-noise conditions by varying the standard deviation of the Gaussian distribution (15 or 30 deg; Fig. 6a). These two conditions were presented in separate blocks, with block order counterbalanced across participants. Within each block, stimulus ensembles of different set sizes were randomly interleaved.

**Figure 6.**
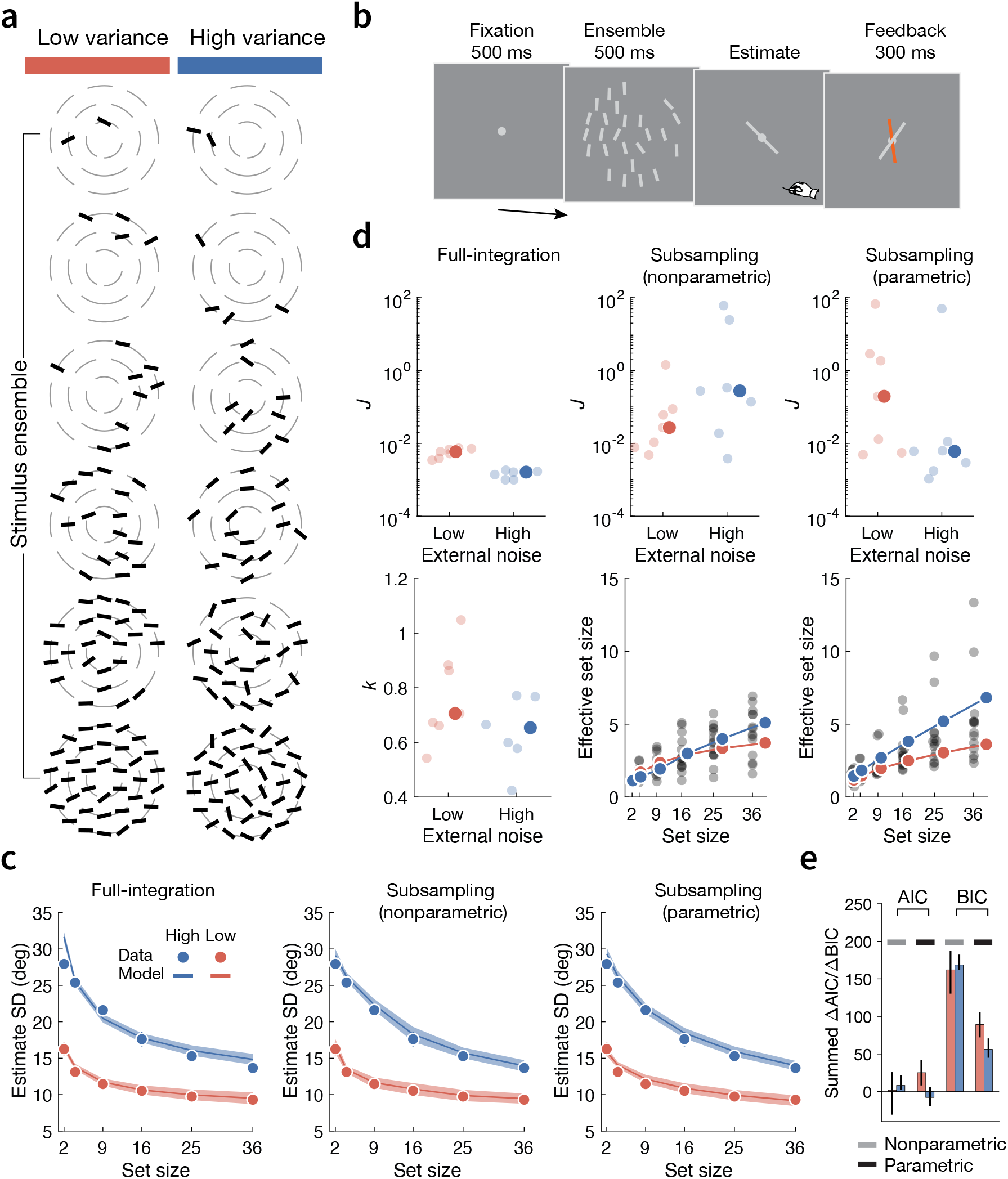
Model fits to data from the ensemble-orientation averaging experiment. (**a**) Example stimulus ensembles. (**b**) Ensemble mean-orientation estimation task. (**c**) Measured and model-predicted SDs of ensemble-mean estimates as a function of set size under low (red) and high (blue) external-noise conditions. Error bars and shaded regions indicate SEMs of the measured data and model predictions, respectively. (**d**) Fitted model parameters. The top row shows the fitted baseline resource parameter *J* for the full-integration (first column), nonparametric subsampling (second), and parametric subsampling (third) models. The bottom row shows the fitted scaling parameter *k* for the full-integration model (first column) and the recovered effective sample sizes for the nonparametric (second) and parametric (third) subsampling models. Dimmed dots indicate individual-participant estimates, and solid dots indicate the group median for *J* and *k* and the group mean for effective sample size. (**e**) Model comparison. Summed AIC and BIC differences between the full-integration model and each subsampling model for both external-noise conditions. Positive values favor the full-integration model.

**Figure 7.**
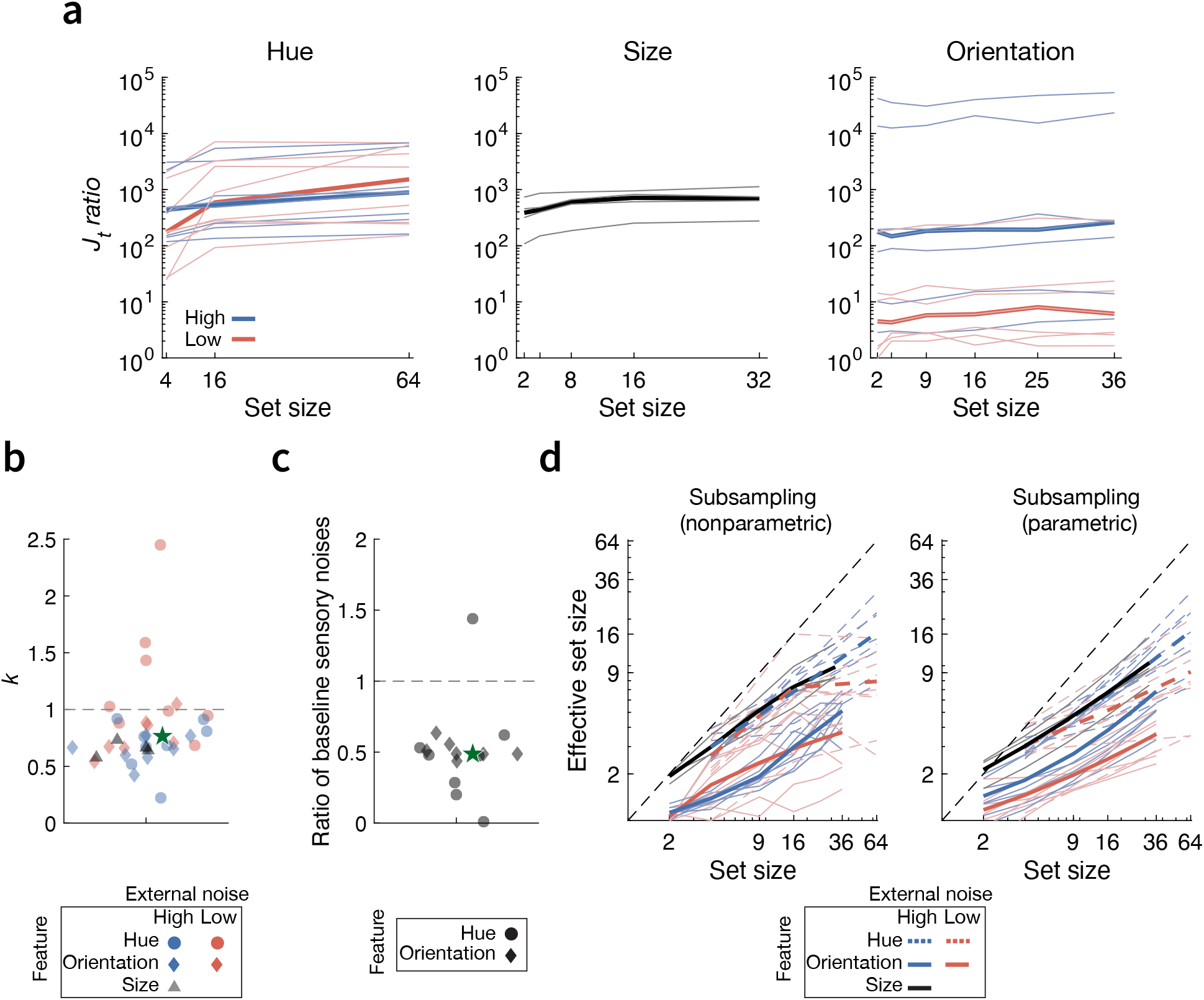
Comparisons of fitted parameters across datasets. (**a**) Ratio of total resource expenditure between the nonparametric subsampling and full-integration models, computed from their fitted parameters for the ensemble-hue (left), ensemble-size (middle), and ensemble-orientation (right) datasets. Thin lines indicate individual-participant ratios, and solid lines indicate the group median. (**b**) Fitted power-law scaling parameter *k* for the ensemble-hue (circles), ensemble-orientation (diamonds), and ensemble-size (triangles) averaging datasets. (**c**) Ratio of baseline sensory encoding noise, *σ*_sens_, between the low and high external-noise conditions for the ensemble-hue and ensemble-orientation datasets, converted from the fitted *J* values. In (**b**) and (**c**), dimmed symbols indicate individual-participant estimates and the green star indicates the group median. (**d**) Recovered effective sample size as a function of physical set size for the ensemble-hue (colored dashed lines), ensemble-orientation (colored solid lines), and ensemble-size (black lines) datasets under the nonparametric (left) and parametric (right) subsampling models. Thin lines indicate individual-participant estimates and thick lines indicate group means. Red and blue denote low and high external-noise conditions, respectively.

Each trial began with a fixation dot presented for 800 ms, followed by the stimulus ensemble for 500 ms. Seven participants were instructed to reproduce the generative mean of the ensemble orientations by moving a probe line with mouse. Veridical feedback was provided after each response to facilitate learning of the ensemble statistics (Fig. 6b). The whole experiment consisted of 1440 trials, equally divided into 6 sessions. The study adhered to the Declaration of Helsinki and was approved by the Institutional Review Board of New York University.

As in the ensemble-hue averaging dataset, we fit the models separately to the two external-noise conditions. Within each condition, each model was fit jointly to the trial-by-trial estimates across all set sizes, separately for each participant. Consistent with the previous two datasets, ensemble-averaging performance improved with increasing set size: the standard deviation (SD) of ensemble-mean estimates decreased as set size increased in both external-noise conditions (Fig. 6c). Predictions from all models again closely tracked the measured SD as a function of set size (Fig. 6c, shaded regions; see Fig A3 for model fits to data of individual participants). The fitted baseline resource parameter *J* values were smaller under the full-integration model than under either subsampling model in both external-noise conditions. Under the full-integration model, the fitted scaling parameter *k* values were below 1 in both conditions, indicating that total coding-resource allocation increased with set size. Under both subsampling models, the recovered effective sample sizes were again substantially smaller than the physical set sizes and were generally smaller in the low-external-noise condition, consistent with the pattern observed in the ensemble hue averaging dataset (Fig. 6d).

To achieve roughly the same ensemble coding accuracy, the full-integration model still required much less total resource expenditure than the non-parametric subsampling model (Fig. 7a, right). Model-comparison results yielded a similar conclusion as before. Once model complexity was taken into account, the subsampling models provided no superior account of the data relative to the full-integration model. AIC values were comparable across models in both external-noise conditions, whereas BIC consistently favored the full-integration model (Fig. 6e).

### Comparison across datasets

Across all three datasets, we consistently find that the full-integration model provides an equally effective, yet more parsimonious, account of ensemble-coding behavior across set sizes, stimulus features, and tasks. Comparing the fitted parameters across datasets further reveals several consistent patterns.

First, under the full-integration model, the fitted power-law scaling parameter *k* is predominantly below 1 (Fig. 7b), indicating that total processing-resource allocation tends to increase with set size. The increase in total resource allocation with set size also aligns with findings from other perceptual and working-memory tasks, in which the inferred power-law scaling parameter *k* is often smaller than 1 (Bays & Husain, 2008; Keshvari et al., 2013; Ma & Huang, 2009). This pattern suggests that the visual system can flexibly adjust its resource investment according to information-processing load: as the amount of visual information increases with set size, allocating additional resources can improve task performance. Such a flexible resource deployment does not contradict the claim that total processing resources are limited. It argues only against the stronger assumption of a rigid visual system that invests a fixed amount of its limited resource budget regardless of visual input load or current task demands.

Second, the fitted baseline coding-resource parameter *J* values under the full-integration model are overall larger in the low external-noise condition than in the high external-noise condition, a pattern observed in both ensemble-hue and ensemble-orientation datasets. Equivalently, the ratio of baseline sensory encoding noise *σ*_sens_, converted from *J* values, between the low and and high external-noise conditions predominantly falls below 1 (Fig. 7c). This difference could reflect a genuine change in baseline coding-resource availability, and hence in item-level sensory encoding precision, across the two conditions; alternatively, it may partly capture differences arising from downstream computations. Because our minimal models deliberately omit downstream sources of variability, for good reasons discussed above, the baseline sensory encoding noise may also absorb variability arising from later computational stages. One possibility, suggested by our on-going modeling work, is that when external stimulus uncertainty is high, downstream processing could discount the sensory evidence to lower computational costs. Such attenuation of sensory signal strength could manifest as reduced “effective” encoding precision.

Lastly, the inferred effective sample size under both subsampling models is strictly smaller than, but also tends to increase with, the actual set size (Fig. 7d), a pattern consistently shown under previous subsampling models. However, the rate of change in effective sample size as a function of actual set size varies considerably across stimulus features, external noise levels, and participants. For instance, under the parametric subsampling model, the fitted power-law scaling parameter *α* (see Eq. 14) ranges from 0.08, indicating that the effective sample size remains nearly constant as physical set size increases, to 1, indicating a linear increase in effective sample size with physical set size (Fig. 7d, right panel). The relation between physical and effective sample size is even more diverse under the non-parametric sampling model (Fig. 7d, left panel). Instead of treating effective sample size as an ad hoc estimated quantity, subsampling accounts require a principled explanation of how effective sample size should scale with physical set size.

## Discussion

In this article, we examined the subsampling hypothesis in ensemble coding and showed that it is neither more resource-efficient nor more effective in accounting for ensemble-coding behavior than a resource-rational full-integration account. Through mathematical analysis, we established that an observer adopting a subsampling strategy must allocate more processing resources than a resource-rational full-integration observer to achieve the same integration precision whenever more than one item is involved. We further demonstrated, through systematic model evaluations across multiple datasets, that subsampling does not provide a superior account of ensemble-averaging behavior across various stimulus features. Model comparison results consistently favored the more parsimonious full-integration account, which achieved comparable explanatory power with fewer free parameters. Our results collectively suggest that there is no theoretical or explanatory advantage of subsampling in ensemble coding.

Instead, we advocate a resource-constrained full-integration account of ensemble coding. Relative to subsampling, full integration provides a more parsimonious account at both the algorithmic and implementation levels. At the algorithmic level, it specifies a common computational strategy for extracting summary statistics across stimulus features and tasks, and offers a principled account of set-size effects without requiring separate, ad hoc estimates of effective sample size for each set size, as is often the case in subsampling models. At the implementation level, full integration can be readily instantiated by population-response models in which neural responses to all ensemble items are pooled, without requiring additional assumptions about the neural mechanisms that selectively exclude items from encoding or downstream readout (Robinson & Brady, 2023; Utochkin et al., 2023). The explanatory scope of the full-integration account also extends beyond set-size effects. Recent work has shown that optimal full-integration models constrained by efficient sensory encoding can quantitatively account for the systematic overweighting of inlier stimuli (i.e., ensemble items closer to a decision reference) in ensemble coding (Ni & Stocker, 2023, 2024, 2026). Given its parsimony and generality, we argue that resource-constrained full integration should serve as the default framework for modeling ensemble-coding behavior.

Our resource-constrained full-integration account does not assume that the total amount of processing resources remains fixed across set sizes. Instead, total resource allocation may increase or, under some conditions, decrease as visual information load changes. This assumption is consistent with evidence that the visual system flexibly adjusts the deployment of limited resources, such as attention and working memory, according to processing demands (Bays et al., 2009; Bays & Husain, 2008; Culham, Cavanagh, & Kanwisher, 2001; Franconeri, Alvarez, & Cavanagh, 2013; Van den Berg et al., 2012), and can rapidly redistribute these resources to prioritize behaviorally relevant information and maximize task performance (Emrich, Lockhart, & Al-Aidroos, 2017; Ni & Stocker, 2023; Yoo, Klyszejko, Curtis, & Ma, 2018). Following previous work, we characterize the relationship between set size and total resource allocation using a power-law scaling function. This choice is primarily descriptive: our theoretical conclusions are largely agnostic about the precise functional relationship between set size and total coding-resource investment, and the power law provides a simple and flexible approximation. Indeed, recent resource-rational analyses suggest that this relationship may be more complex, with total resource investment varying non-monotonically with set size as a consequence of an optimal tradeoff between behavioral performance and the neural cost of resource expenditure (Van den Berg & Ma, 2018). From this perspective, the power-law scaling used here can be viewed as a tractable approximation to a potentially more complex resource-allocation function rather than as a fundamental law governing resource deployment. Identifying the optimal relationship between resource allocation and set size in ensemble coding would be an important direction for future work.

The full-integration account does not require limited coding resources, such as attention, to be distributed equally across all ensemble items. Instead, the visual system appears capable of flexibly reallocating its limited resources toward particular items within an ensemble. As a result, items that receive greater coding resources and are therefore encoded with higher precision, such as those pre- or post-cued (Choi & Chong, 2020; De Fockert & Marchant, 2008; B. Li, Wang, Zhang, & Qian, 2024), physically more salient (Iakovlev, Khvostov, Ásgeirsson, Utochkin, & Kristjánsson, 2025; Iakovlev & Utochkin, 2021; Im, Park, & Chong, 2015; Kanaya, Hayashi, & Whitney, 2018), displayed in the central visual field (Dandan, Ji, Song, & Sayim, 2023; Ji, Chen, & Fu, 2014; Pascucci, Ruethemann, & Plomp, 2021; Tiurina, Markov, Whitney, & Pascucci, 2024), and associated with high probabilities (de Gardelle & Summerfield, 2011; V. Li, Castañón, Solomon, Vandormael, & Summerfield, 2017; Ni & Stocker, 2023, 2024, 2026), contribute more strongly to the resulting summary representation. This form of precision-weighted averaging can be readily incorporated into optimal full-integration models (Ni & Stocker, 2023, 2026).

Lastly, although we argue here against subsampling of stimulus-feature representations in ensemble coding, our results do not necessarily challenge other forms of sampling limitation in vision. One important distinction is between feature-level subsampling and spatial sampling that occurs at early stages of visual processing. Because the retinal and early visual systems have finite spatial sampling densities, the retinal image cannot be represented with arbitrarily high spatial resolution. In particular, the density of retinal ganglion-cell receptive fields decreases substantially with eccentricity, imposing progressively stronger limits on spatial resolution in peripheral vision (Barlow, 1979; Watson, 2014). Spatial sampling thus concerns the resolution with which the visual image itself is represented: insufficient sampling density limits how much spatial information can be recovered from the input. Feature-level subsampling in ensemble coding, by contrast, concerns which subset of spatially resolved local feature representations is incorporated into the computation of summary statistics. In most ensemble-averaging paradigms, the individual elements are suprathreshold, spatially separated stimuli with discernible feature values. The subsampling hypothesis in the context of ensemble coding is thus not simply that early vision has finite spatial resolution, but that only a subset of these available feature representations is selected for integration. Our analyses do not question spatial sampling or the use of related approaches, such as equivalent-noise methods, for characterizing its efficiency (e.g., Zeevi & Mangoubi, 1984). Rather, they call into question feature-level subsampling accounts in which ensemble feature statistics are assumed to be computed from only a discrete subset of otherwise available stimulus-feature representations.

## Conclusion

The human visual system continuously compresses a vast stream of sensory information into summary representations, reducing processing demands under limited resources. Our analyses suggest that such compression is better explained by integrating information across the full ensemble rather than by selectively sampling only a subset of items. Compared with subsampling, resource-constrained full integration achieves the same level of ensemble-coding accuracy with substantially lower resource expenditure and provides a more parsimonious, yet equally effective, account of behavior across feature domains. We therefore conclude that there is no benefit of invoking the subsampling hypothesis in explaining ensemble coding behavior.

## Acknowledgment

We thank Lari S. Virtanen for sharing the data and Baek and Chong (2020) for making their dataset publicly available. This research was supported by NIH grant EY08266 and the NYUAD Center for Brain and Health, funded by Tamkeen under NYU Abu Dhabi Research Institute grant CG012.

## APPENDIX A

Here we derive the posterior of the generative ensemble mean *µ* given the sensory measurement set *X, p*(*µ* | *X*). We first compute the likelihood for one sensory measurement *x*_*i*_

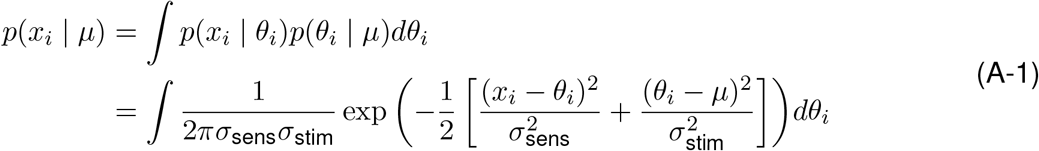

Expanding the quadratic inside the exponential:

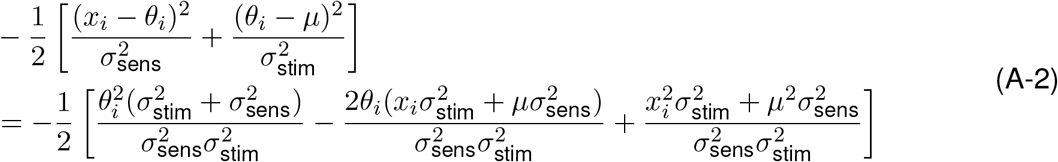

Let

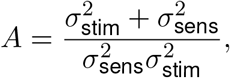

and

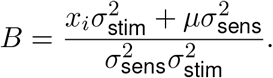

Substituting and completing the square, we have:

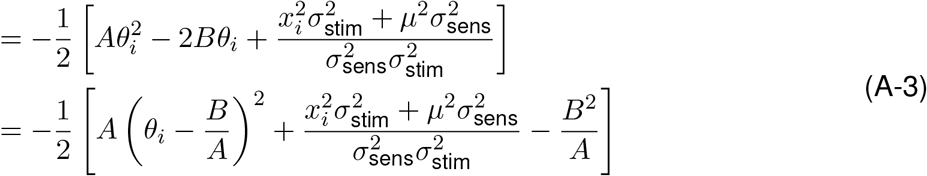

Back to Eq. A-1,

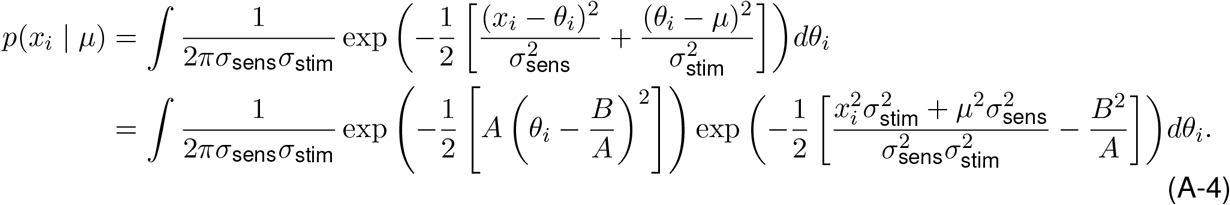

To compute the integral

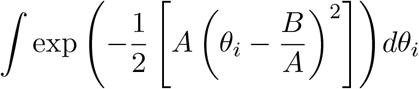

Let

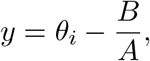

then

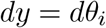

The integral over *θ*_*i*_ then becomes

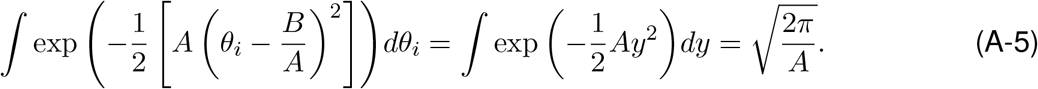

This is because the (definite) integral for a Gaussian function exp(−*ay*^2^) is 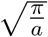 for *a >* 0.

Back to the main calculation,

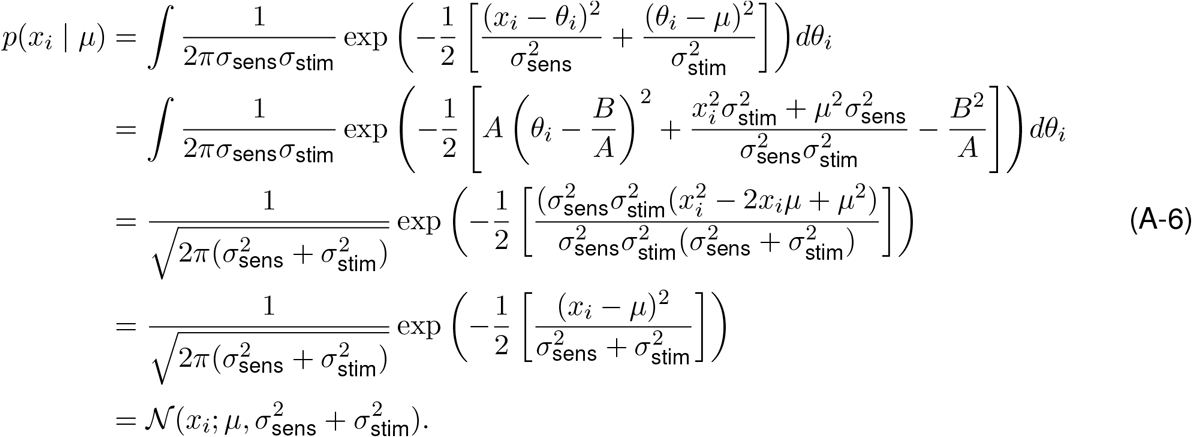

The posterior distribution *µ* given the entire sensory measurement set *X* is given as follows:

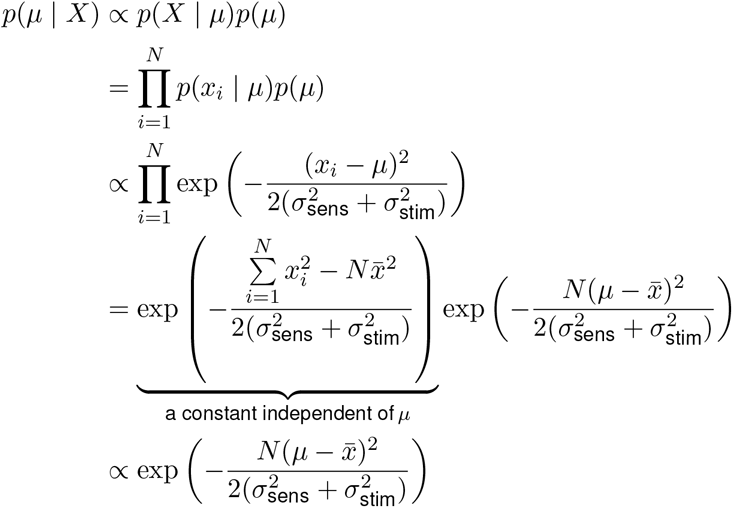

Re-normalizing this Gaussian function by its integral 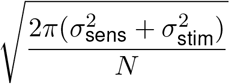 (see Eq. A-5), we have:

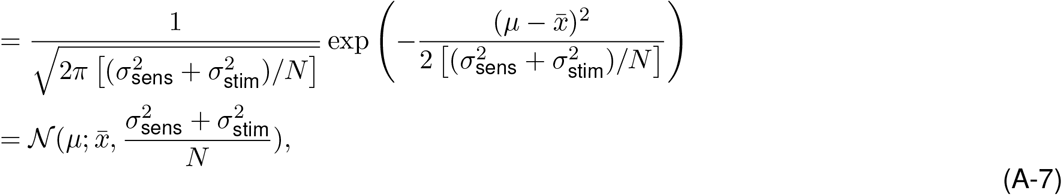

in which the prior *p*(*µ*) is assumed to be uniform and *N* is the ensemble size.

## Appendix B

In this section, we derive the distribution of the ensemble-mean 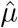 estimates given one particular stimulus ensemble Θ. Under the Gaussian-noise assumption, it is computed as follows:

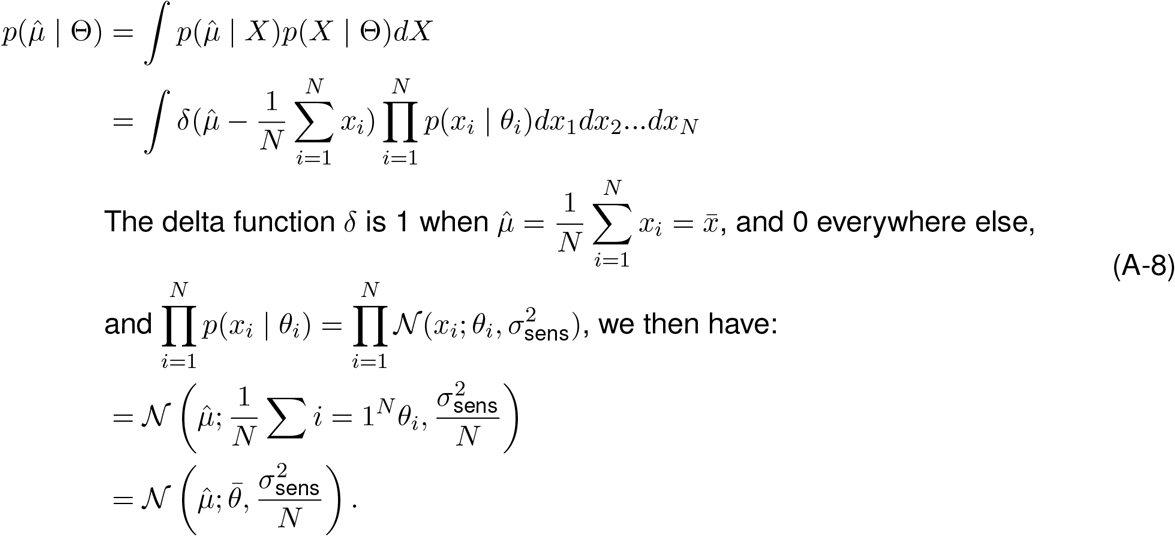

## Appendix C

Our main conclusion that full integration is more resource-efficient holds when the stimuli are instead sampled from a uniform distribution with a fixed range and varying mean. Specifically, we assume that each stimulus *θ*_*i*_ is independently drawn from a uniform distribution centered on a generative *µ* with half-range *h*:

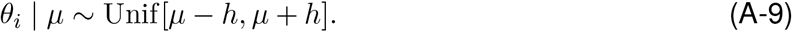

The sensory measurement of each stimulus is assumed to be corrupted by independent Gaussian noise:

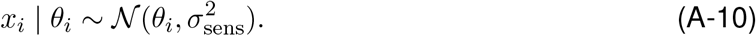

Under these assumptions, the posterior over the generative *µ* is

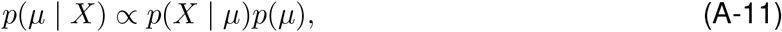

where *X* = (*x*_1_, …, *x*_*N*_) and *p*(*µ*) is assumed to be a flat prior. The likelihood factorizes across items:

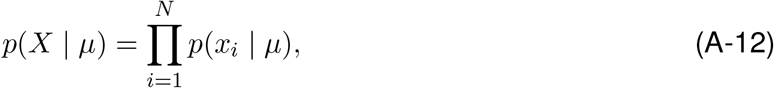

with

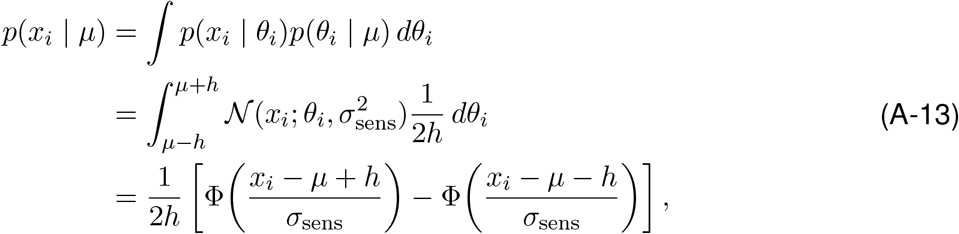

where Φ(·) denotes the cumulative distribution function of the standard normal distribution. Therefore, we have

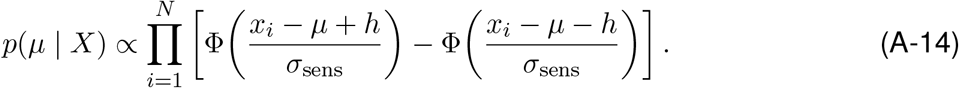

Unlike the Gaussian generative case, the posterior *p*(*µ* | *X*) is generally not Gaussian.

We next compute the distribution of the mean estimate conditional on the realized stimulus ensemble 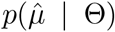, where Θ = (*θ*_1_, …, *θ*_*N*_). Conditional on Θ, the sensory measurements are independent:

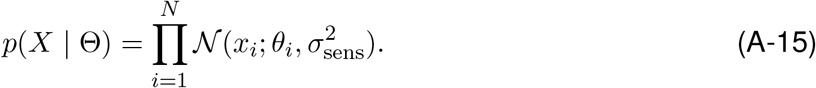

Because 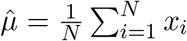 is a linear combination of independent Gaussian random variables, 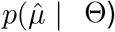 is also Gaussian, with its mean

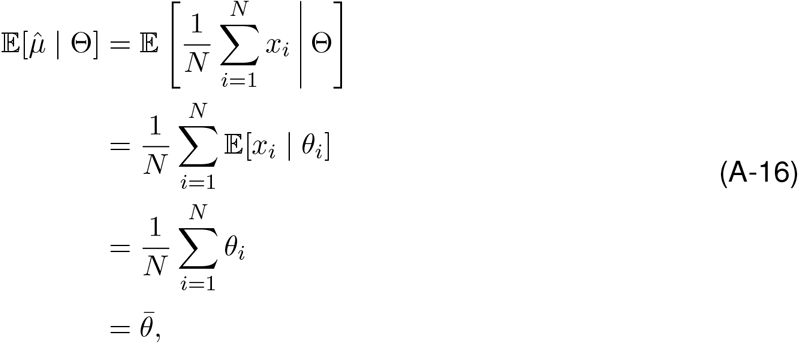

and its variance

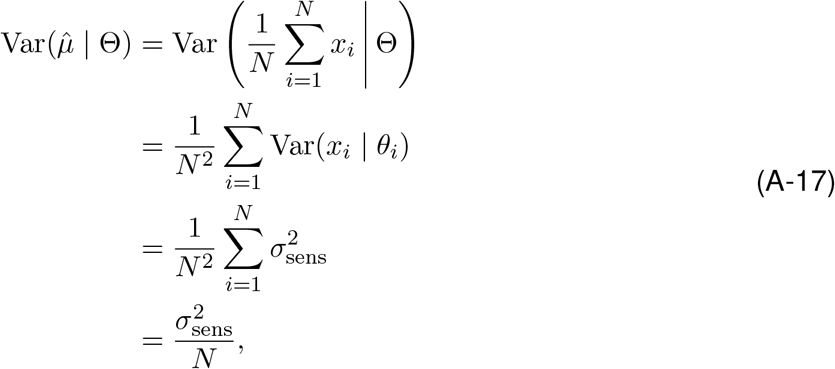

where the cross-covariance terms vanish because the sensory measurements are conditionally independent given Θ. Therefore,

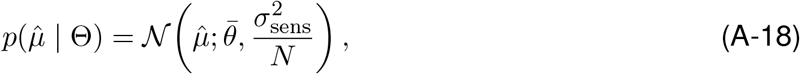

which is the same as in the Gaussian case.

Finally, we can compute the distribution of the mean estimate conditional on the generative mean, 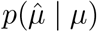 by marginalizing over the realized stimulus set:

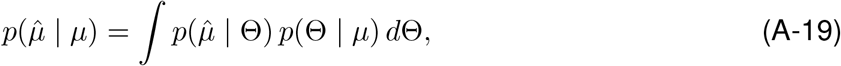

where

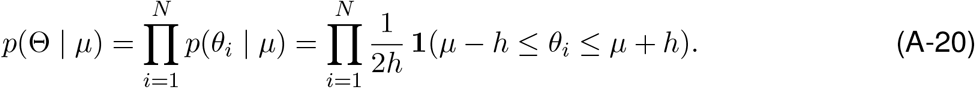

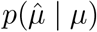 is no longer Gaussian in general, but we can derive its mean and variance. Since

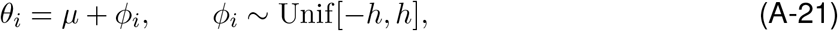

we have

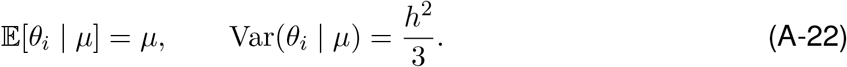

Given Gaussian sensory noise,

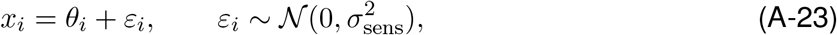

we have

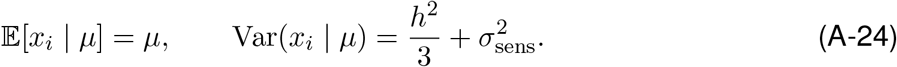

Therefore, for the mean estimate

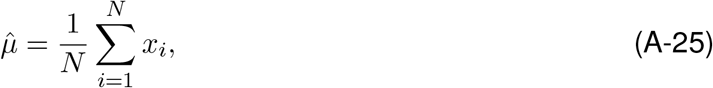

we obtain

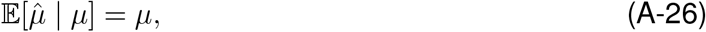

and

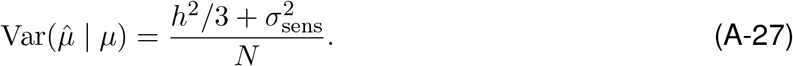

### Subsampling versus full integration

We now compare the total encoding resource required by a subsampling strategy and a full-integration strategy. Assume that under full integration, the encoding precision allocated to each item is

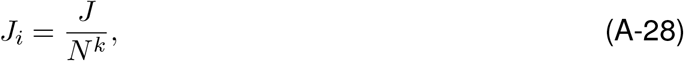

so that the corresponding sensory noise variance is

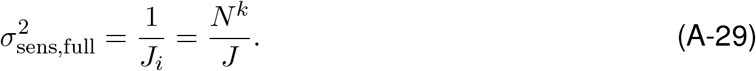

Substituting into Eq.A-27, the variance of the mean estimate under full-integration account then becomes

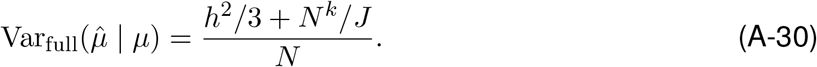

Under a subsampling strategy, only *M* items are sampled, each encoded with baseline precision *J*, so that

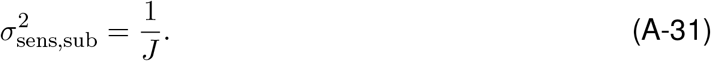

Then the variance of the mean estimate is

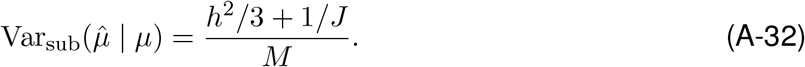

To achieve the same performance, the two variances must be equal:

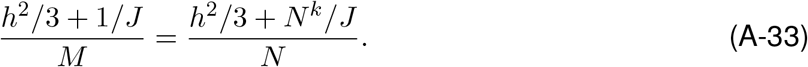

Solving for *M* yields

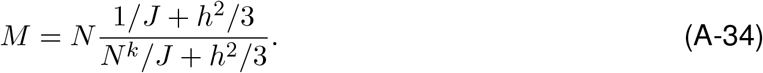

The total resource required by the subsampling strategy is

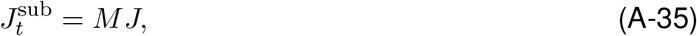

whereas the total resource required by the full-integration strategy is

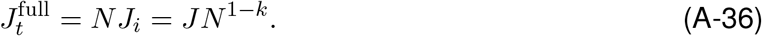

Their ratio is therefore

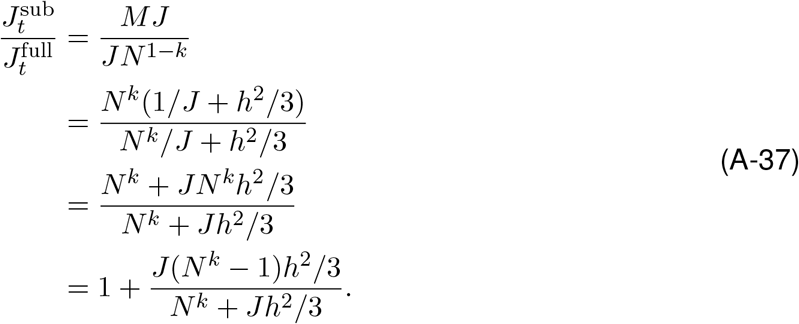

Hence,

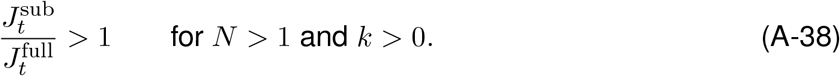

Thus, under the uniform generative model, subsampling requires more total encoding resource than full integration to achieve the same estimation performance. Equality holds when *k* = 0, and the inequality reverses when *k <* 0.

## Appendix D

In this section, we show that the main conclusion still holds when the observer instead estimates the sample mean,

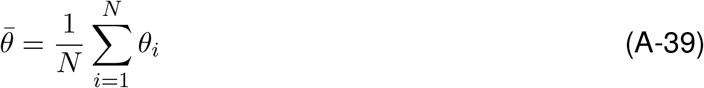

rather than the latent generative mean *µ*. Assuming a flat prior on 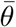 and independent Gaussian sensory noise, the posterior over the sample mean is

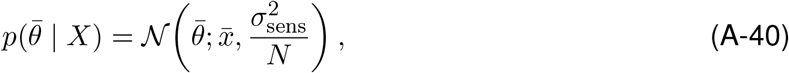

in which 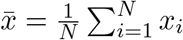.

The distribution of the sample mean estimate 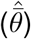 given a realized stimulus set (Θ) is also Gaussian

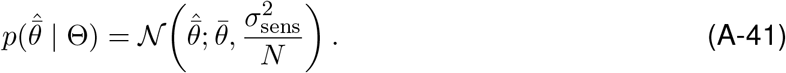

The sensory encoding noise for each of the *N* items is

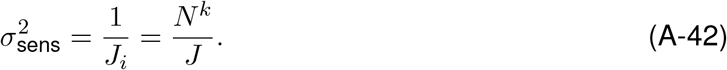

The estimate variance relative to the sample mean becomes

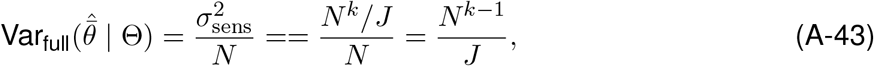

and the total coding resource expenditure is

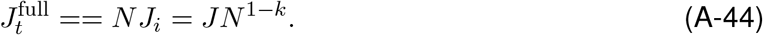

### Subsampling

Under subsampling, the observer samples only *M* out of the *N* items, and encodes each sampled item with baseline precision *J*, so that

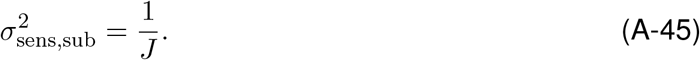

We define *S* as a subset of *M* sampled items, and define the estimate of this subset

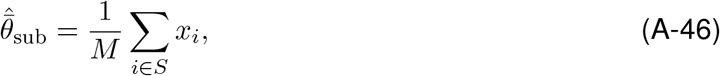

and the actual mean of this subset is

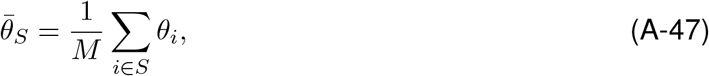

The variance of estimates relative to the sample mean can be decomposed into two components:

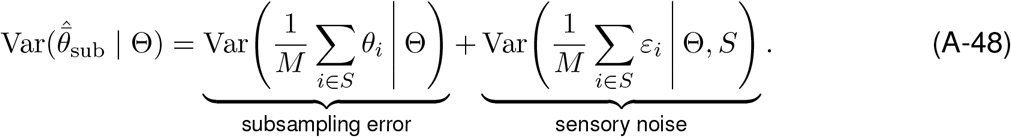

The first component is due to sampling noise (i.e., which *M* items are sampled), and the second component is caused by sensory encoding noise.

The sensory-noise part is computed as

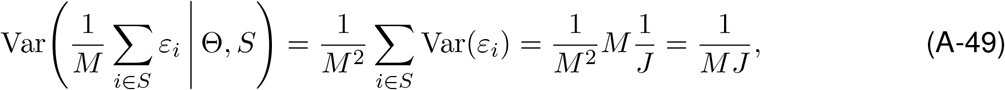

in which *ε*_*i*_ is the independent sensory noise for each stimulus, 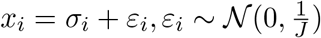.

The sampling error component is computed as

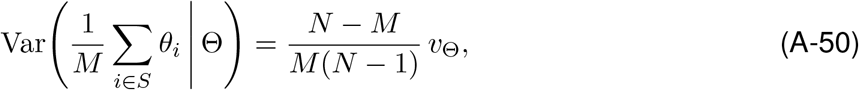

in which *v*_Θ_ is the population variance of a given stimulus set

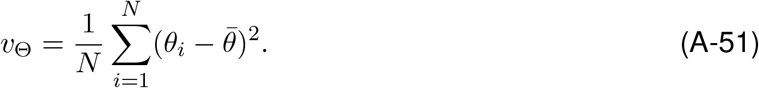

Therefore, the total variance becomes

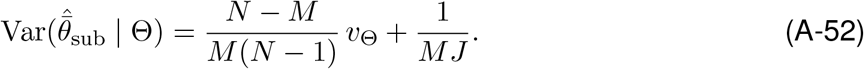

### Gaussian generative distribution

We first consider the case in which the stimuli are independently sampled from a Gaussian distribution,

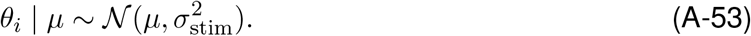

Then the expected value of *v*_Θ_ is

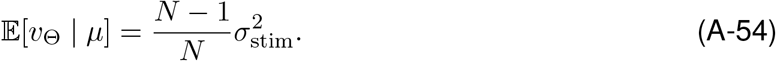

Substituting in Eq. A-52, we have

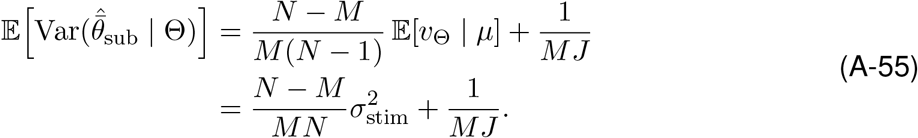

To match the performance under the full-integation and subsampling models, we have:

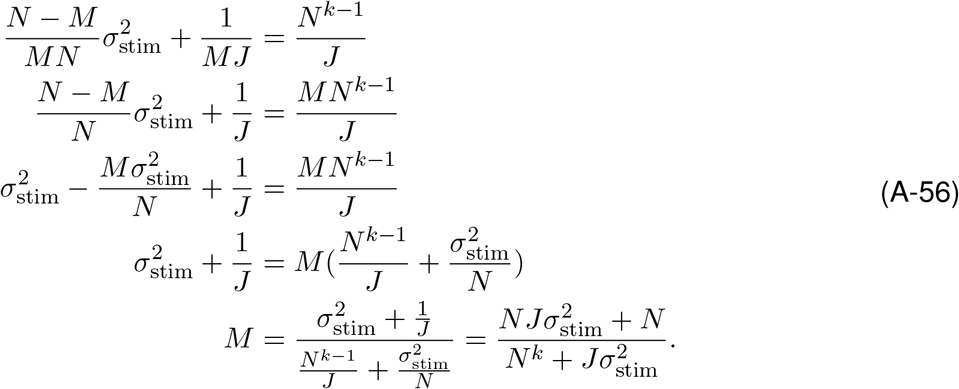

The corresponding total resource under subsampling then becomes

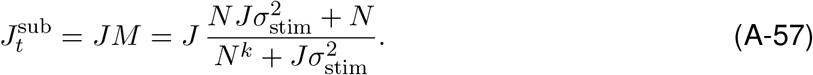

The ratio of total resource expenditure under the two models is

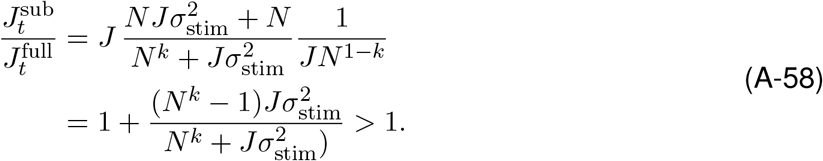

The above equation holds true as long as *k >* 0 and *N >* 1.

Thus, when observers estimate the sample mean instead, subsampling is still always less resource-efficient than full integration.

### Uniform generative distribution

Now consider the case in which the stimuli are sampled from a uniform distribution with fixed half-range *h*,

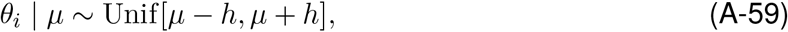

The variance of this distribution 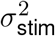 then becomes

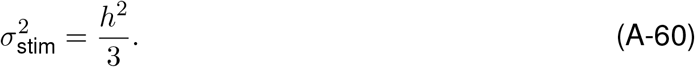

Plugging it into Eq. A-55, we have

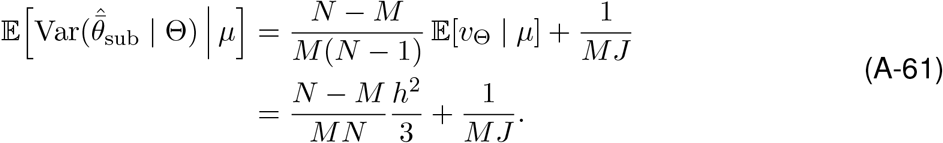

Equating this with the full-integration variance gives

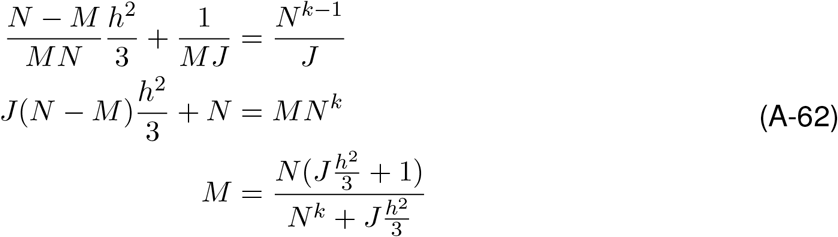

The total resource ratio becomes

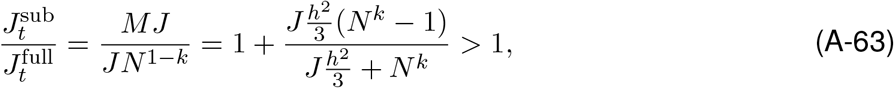

whenever *k >* 0 and *N >* 1. Thus, the same qualitative conclusion holds for the uniform generative case.

## Appendix E

**Figure A1.**
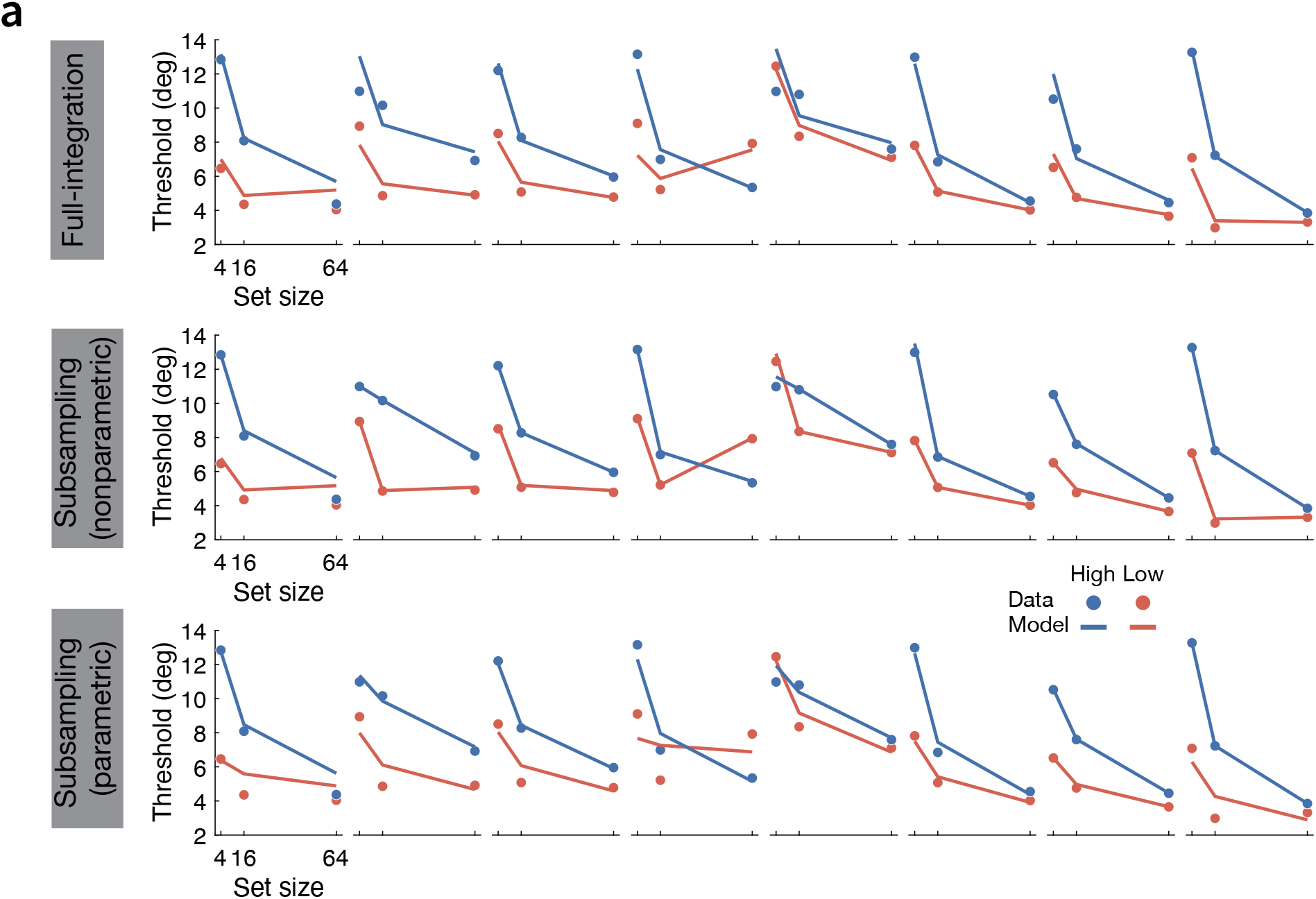
Model fits to individual-participant data from the ensemble-hue averaging experiment of Virtanen et al. (2020). (**a**) Measured and model-predicted thresholds of discriminating ensemble mean hue as a function of set size under low (red) and high (blue) external-noise conditions for each of the eight participants (columns), shown for the full-integration model (first row), nonparametric subsampling model (second row), and parametric subsampling model (third row).

**Figure A2.**
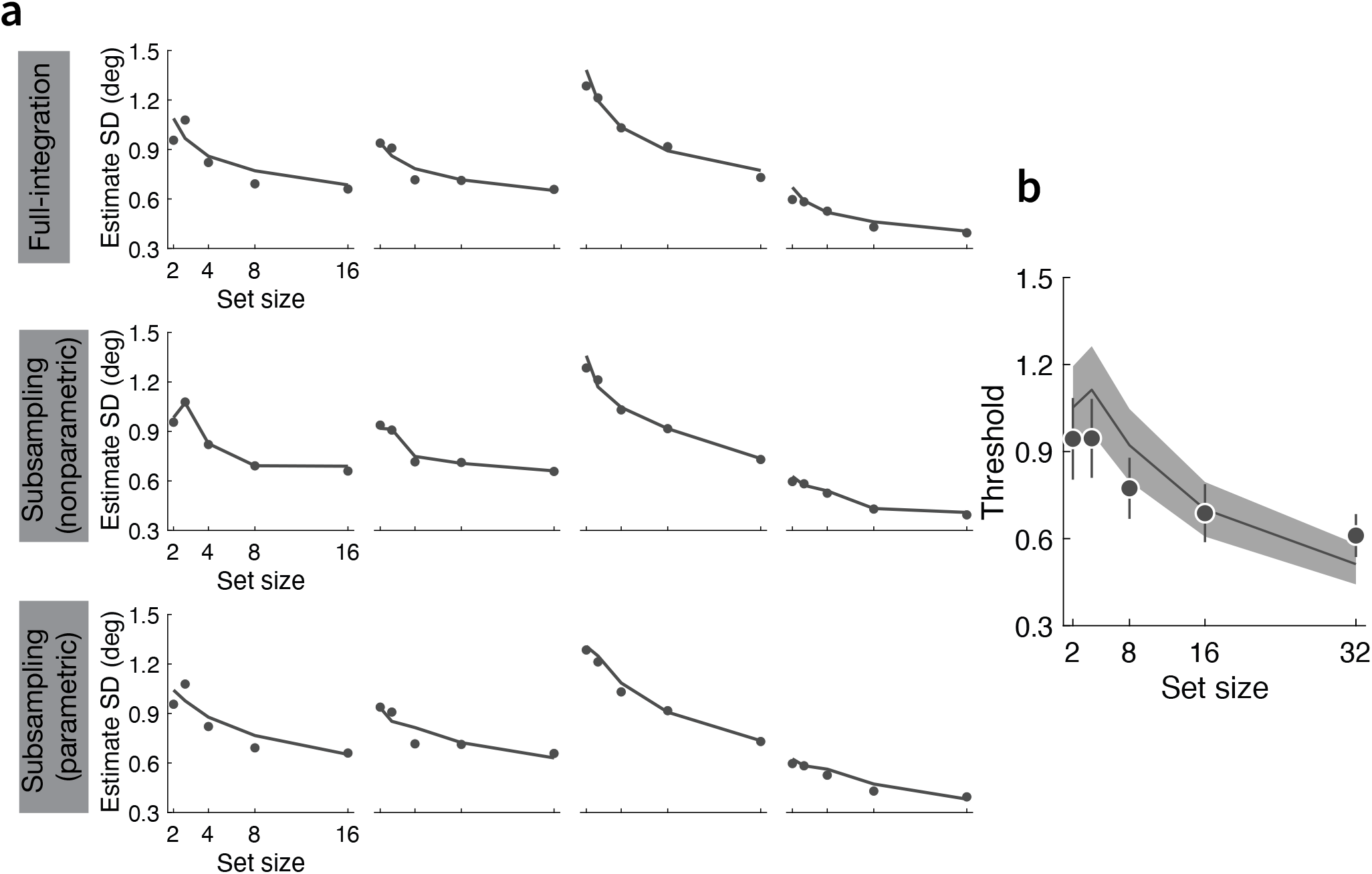
Model fits to individual-participant data from the ensemble-size averaging experiment of Baek and Chong (2020). (**a**) Measured and model-predicted thresholds of discriminating ensemble mean size as a function of set size for each of the four participants (columns), shown for the full-integration model (first row), nonparametric subsampling model (second row), and parametric subsampling model (third row). (**b**) Group-average fit of the heuristic full-integration model proposed in Baek and Chong (2020). This heuristic model has the same number of parameters (*J* and *A*) as our full-integration model, but performs consistently worse than ours for all four participants (Δlog-likelihood. 13.3±3.58 (mean ± SEM)).

**Figure A3.**
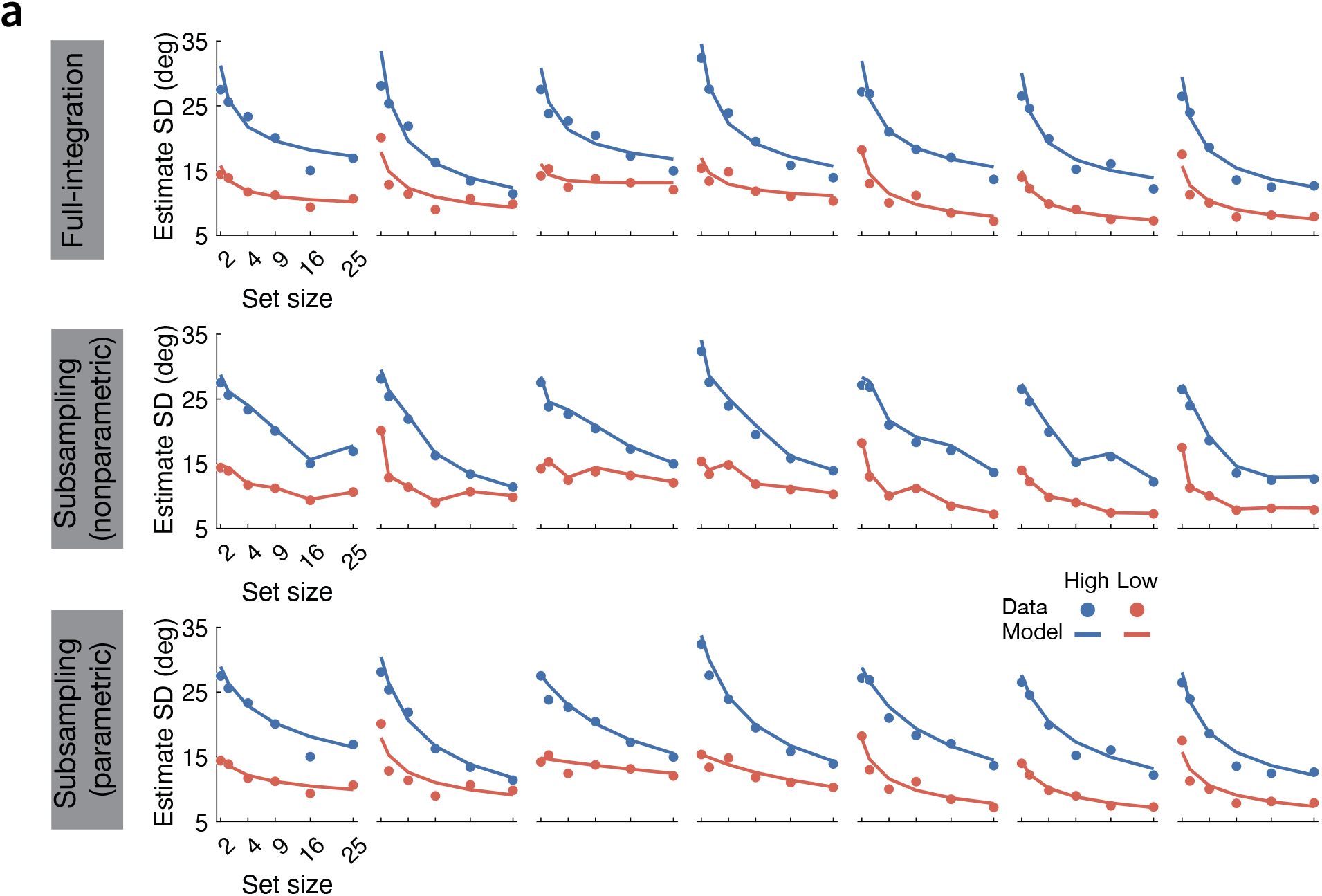
Model fits to individual-participant data from the ensemble-orientation averaging experiment. (**a**) Measured and model-predicted SDs of ensemble-mean estimates as a function of set size under low (red) and high (blue) external-noise conditions for each of the eight participants (columns), shown for the full-integration model (first row), nonparametric subsampling model (second row), and parametric subsampling model (third row).

## Footnotes

1 In the zoom-lens model, sensory encoding noise is given by 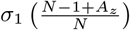, where *σ*_1_ denotes baseline encoding noise, *N* is set size, and *A*_*z*_ is a free parameter governing the attentional modulation effect. The motivation for this particular functional form was not specified in their paper.

